# Mechanosensitive and Reversible Chromatin-Lamina Dewetting Triggers Cellular Contraction During Wound Healing

**DOI:** 10.1101/2025.09.10.674927

**Authors:** Guillermo Bengoetxea, Lenka Backová, Ignacio Arganda-Carreras, Monika Elzbieta Dolega, Ion Andreu, Jérôme Solon

**Author notes:** These authors contributed equally to the work (Guillermo Bengoetxea, Lenka Backová).

## Abstract

Tissue wounding causes rapid mechanical changes in the epithelium, but how these changes affect intracellular organization remains unclear. Using the *Drosophila* embryonic epidermis, we show that wounding induces a fast, transient compaction of chromatin and its detachment (“dewetting”) from the nuclear lamina in about half of the wound-edge cells. Within minutes, chromatin in these cells re-expands and re-associates with the lamina (“re-wetting”). This reversible chromatin–lamina dewetting is mechanosensitive, requiring both the LINC complex and intracellular calcium. During compaction, calcium is stored within chromatin and is later released during ATR-mediated expansion and lamina re-wetting. Chromatin re-expansion then triggers actomyosin contraction and promotes tissue repair through Jun kinase activation, straightening of the wound edge, and supracellular actin ring formation. Together, our findings reveal a multiscale mechanosensitive mechanism by which tissue-scale mechanical changes induce a reversible chromatin-lamina dewetting, leading to cellular contraction and initiation of tissue repair.

## Introduction

Epithelial tissues experience diverse mechanical perturbations throughout the life of a living organism. Such perturbations can generate wounds, disrupting epithelial integrity and triggering rapid mechanical and chemical changes (*1*). Injury initiates a calcium wave that propagate across the tissue via ion channels and sequential ER-stored calcium release, initiating repair programs and immune cell recruitment (*2–5*). Concurrently, the actomyosin cytoskeleton organizes around the wound to generate contractile forces that promote the sealing of the epithelial gap (*6–8*). Several signaling pathways are activated in cells neighboring the wound, including the Jun kinase (JnK) cascade (*1*), which is essential for actomyosin reorganization and force generation at the wound leading-edge (*9–11*). Mechanosensitive activation of tissue repair machinery has been reported, with potential involvement of pathways such as Hippo, mechanosensitive channels like Piezo, and nuclear mechanosensing (*12–14*). Notably, blocking yes-associated protein (YAP) post-injury can reprogram wound healing toward scar-free regeneration (*15*).

*In vitro* studies have established that cells respond to mechanical stress by activating pathways such as Hippo and reorganizing their cytoskeleton (*16*, *17*). Beyond the cytoplasm, mechanical cues also strongly influence nuclear architecture. Forces transmitted through the cytoskeleton and the LINC complex can directly deform the nucleus and alter chromatin organization, transforming the nucleus from a passive compartment into a mechanosensory hub that integrates external tension with genome function. Nuclear envelope stretching triggers calcium release, driving inflammation (*14*), membrane blebbing (*18*), migration (*19*), and chromatin rheological changes (*20*). The ataxia telangiectasia and Rad3-related (ATR) kinase senses mechanical deformation at the nuclear periphery even in the absence of DNA damage, linking chromatin state to stress checkpoints (*21*) and actin reorganization (*22*).

*In vivo*, chromatin and nuclear mechanosensing also occurs, particularly during tissue injury and inflammation. Rapid or repeated tensional changes remodel chromatin, activating genetic programs that drive repair or scarring (*23*, *24*). These changes in chromatin organization impact nuclear envelope mechanics and subsequently overall cellular mechanics (*23–25*). For instance, nuclear envelope stretching, resulting from osmotic swelling at the wound site, has been shown to initiate inflammatory signaling and immune recruitment (*14*). Mechanosensitive multiscale connections, involving nuclear and chromatin architecture, are clearly central to the response of biological tissues to external stresses. However, the mechanisms underlying the dynamic connectivity and remodeling of nuclear structures during the cellular response remain incompletely understood.

Here, we identify a reversible, mechanosensitive dewetting of chromatin from the nuclear lamina that triggers cellular contraction and promotes tissue repair. Loss of tension during wounding induces, in about half of the wound-edge cells, rapid chromatin compaction and lamina detachment, followed within minutes by ATR-mediated chromatin re-expansion, calcium release, and re-wetting. This transition promotes actomyosin contraction, actin ring formation, and early JnK activation, establishing a pathway by which tissue-scale mechanical changes directly remodel nuclear organization to drive cellular and tissue contractility.

## Results

### Epithelial wound generation triggers rapid chromatin compaction and re-expansion in neighboring cells

To assess the impact of wound generation on chromatin organization and probe nuclear-scale mechanosensing, we performed high-resolution confocal imaging in *Drosophila* embryos expressing mRFP-tagged histones (His2Av-mRFP). Precise wounds of 25 μm length in the anterioposterior (AP) direction were induced in the embryonic epidermis using a pulsed UV laser (**Fig. 1A**, see Methods).

**Figure 1:**
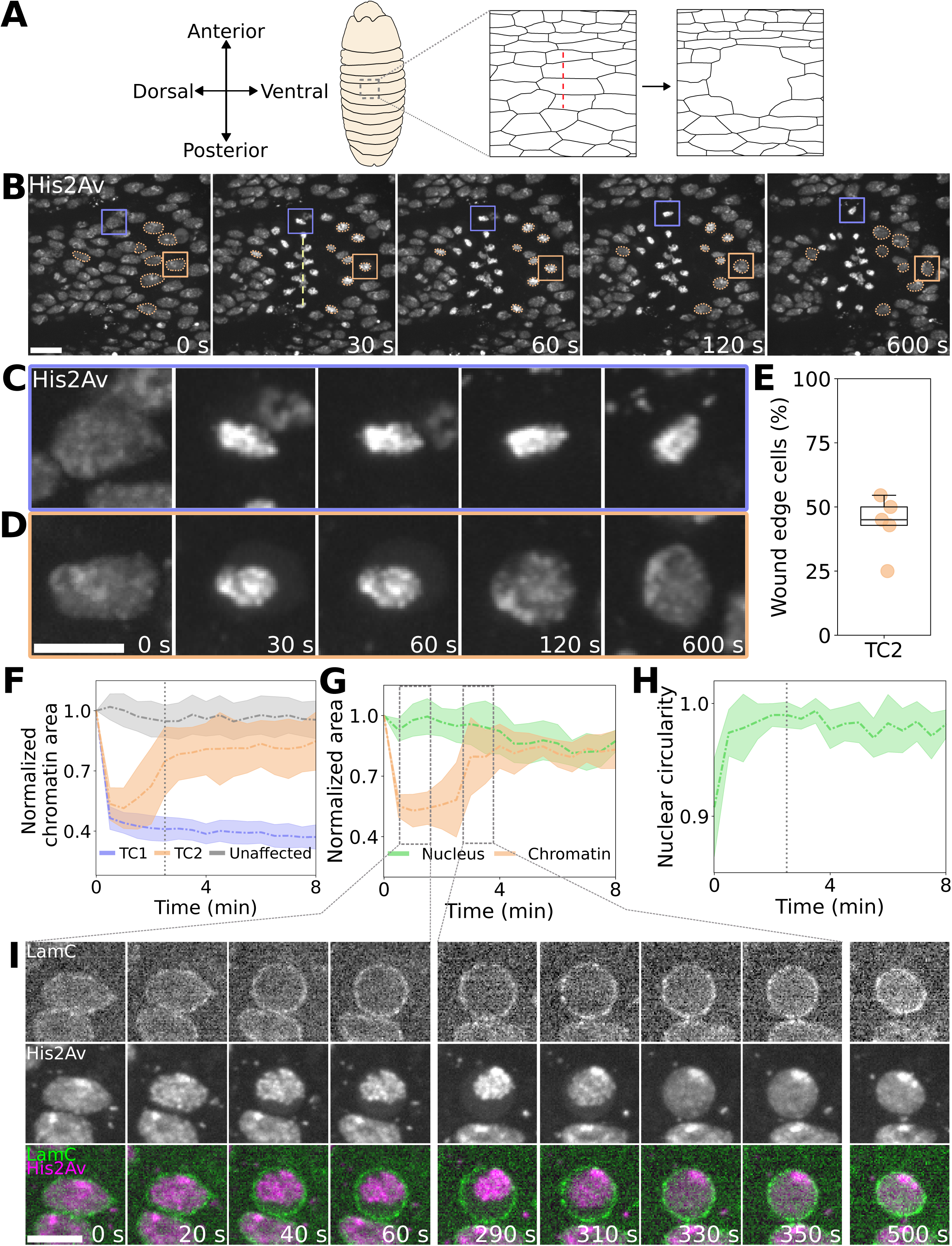
Reversible chromatin-lamina dewetting during wound generation in the Drosophila embryonic epidermis. **(A)** Schematic of the experiment. Wounds of 25 μm in the AP direction are induced in the epidermis of *Drosophila* embryos at late developmental stage. **(B)** Time-lapse images showing the rapid compaction of chromatin in cells neighboring the induced wound (shown in yellow) in an embryo expressing His2Av-mRFP. Two types of behavior can be observed: cells compacting their chromatin without subsequent recovery (TC1, example shown in a blue bounding box), and cells displaying fast, reversible chromatin compaction (TC2, all cells outlined with an orange dashed line). Scale bar 10 μm. **(C)** Time-lapse images showing a representative TC1 cell, which compacts the chromatin without subsequent decompaction. **(D)** Time-lapse images showing a representative TC2 cell, displaying fast, reversible chromatin compaction. Scale bar 5 μm. **(E)** Boxplot of the ratio of wound-edge cells, which undergo reversible compaction (*N* = 5 embryos, *n_TC2_* = 42 cells, *n_WE_* = 98 cells). The black line represents the median of the data, the box encloses 50% of the data around the median, the upper whisker extends from the box to the largest data point lying within 1.5x the inter-quartile range (IQR) from the box. **(F)** Normalized chromatin area of cells displaying the following chromatin behavior: irreversible compaction (TC1 cells in blue), reversible compaction (TC2 cells in orange), and a subset of cells unaffected by the induced wound (in gray) (*N* = 6 embryos, *n_TC1_* = 98 cells, *n_TC2_* = 59 cells, *n_unaffected_* = 100 cells, median + median absolute deviation (mad) shown). **(G)** Normalized area of the chromatin (orange) and the nuclear lamina (green) in the midplane of the nucleus in TC2 cells (*N* = 5 embryos, *n_TC2_* = 23 cells, median + median absolute deviation (mad) shown). **(H)** Graph showing an increase in circularity of the nuclear lamina of TC2 cells (*N* = 5 embryos, *n_TC2_* = 23 cells, median + median absolute deviation (mad) shown). The dashed gray vertical line represents the median decompaction time. After re-expansion, the circularity deviation increases as the nulcear lamina changes shape after the re-wetting. **(I)** Time-lapse showing the chromatin diwetting from the nuclear lamina and the subsequent re-wetting in an embryo expressing Lamin-GFP and His2Av-mRFP. While the chromatin is compacted, the nuclear lamina adopts a near-spherical shape. Subsequently, chromatin re-wet the lamina, leading to a gradual change in the shape of the nucleus. Scale bar 5 μm.

In less than a minute after wound generation, we observed chromatin compaction in most of the cells surrounding the wound (**Fig. 1B-C-D** and **Movie S1**). Surprisingly, in a substantial subset of these cells, chromatin re-expanded after a few minutes (**Fig. 1D, Movie S1** and **Movie S2**). Using nuclear segmentation in embryos co-expressing His2Av-mRFP and sGMCA to label histones and actin, respectively (see Methods), we quantified that ∼50% of all wound-edge cells undergo this reversible compaction (**Fig. 1E**). In these cells, chromatin area shrank by ∼50% for 2–3 minutes before re-expanding to near pre-wound size (**Fig. 1F**). Similar transient chromatin compaction was observed following mechanical wounding of the epidermis, ruling out potential artifacts from UV-induced damage (**Fig. S1A**).

Together, these data reveal two mechanically responsive chromatin behaviors triggered by wounding: (i) a population of cells that compact their chromatin without subsequent decompaction (TC1) (**Fig. 1C**), and (ii) a population displaying fast, reversible chromatin compaction (TC2) (**Fig. 1D-F**).

### Wounding triggers a reversible chromatin-lamina dewetting that restores anisotropic forces on the nucleus

We next examined how the nuclear lamina responds in the population of wound-edge cells that undergo transient chromatin compaction (TC2 cells). Dual imaging of the nuclear lamina (Lamin-GFP) and chromatin (His2Av-mRFP) revealed that during the compaction phase in TC2 cells, chromatin detached from the lamina while nuclear mid-area increased, reassembling onto the nuclear lamina after a few minutes (4.02 ± 3.8 min on average) (**Fig. 1G–I** and **Movie S3**). Such disassembly resembles the dewetting of a polymer from a surface that retracts and minimizes surface contact (*26*). Therefore, this transition can be defined as a transient chromatin-lamina dewetting. TC1 cells remain in a persistent dewetting state and are thus referred to as De-Wet cells, whereas TC2 cells reattach chromatin to the lamina and are termed Re-Wet cells.

High-resolution imaging shows that the nucleus adopts a near-spherical shape during the dewetting phase, indicating a loss of anisotropic forces on the nuclear envelope. This suggests transient disruption of mechanical coupling on both sides of the nucleus, between chromatin and lamina on one side, and cytoskeleton and lamina on the other. Consistently, we observed a significant decrease in cortical F-actin (sGMCA) in Re-Wet cells compared to the surrounding cells in the epidermis (**Fig. S1B**). Within a few minutes, chromatin re-engages the lamina, re-wetting the nuclear envelope. Concomitantly, the nucleus regains an irregular shape with dynamic fluctuations (**Fig. 1G, H, I** and **Movie S3**) and sGMCA levels also return to values comparable to neighboring cells (**Fig. S1B**), indicative of restored force transmission.

Altogether, we identify a reversible chromatin-lamina dewetting process in cells neighboring the wound. This nuclear-scale remodeling distinguishes two mechanically responsive cell populations: De-Wet cells (TC1), which maintain a persistent dewetting state, and Re-Wet cells (TC2), which undergo transient dewetting. This behavior reflects a potential loss and recovery of nuclear–cytoskeletal connectivity in response to tissue-scale mechanical disruption.

### Re-Wet cells maintain compartment integrity during transient chromatin-lamina dewetting

To assess whether transient chromatin-lamina dewetting is associated with damage in plasma membrane or nuclear envelope integrity and subsequent nuclear or cytoplasmic leakage, we expressed either NLS-GFP or freely diffusing GFP in the embryonic epidermis using the UAS-Gal4 system. This revealed two distinct cellular responses matching the previously defined De-Wet cells (TC1) and Re-Wet cells (TC2). The first population consistently shows leakage of both nucleoplasmic and cytoplasmic contents (**Fig. S2A-B-C-D**). These cells fail to reintegrate into the tissue and are subsequently cleared by hemocytes during the healing process. In contrast, Re-Wet cells retain intact nuclear and cytoplasmic compartmentalization throughout the process (**Fig. S2A-B-C-D**) and persisted in the tissue post-healing, indistinguishable from neighboring cells (**Fig. S2E**).

These findings indicate that Re-Wet cells maintain compartment integrity despite transient nuclear remodeling, while persistent dewetting correlates with irreversible damage and cell elimination. Importantly, nuclear and cytoplasmic leakage emerges as an early predictor of whether chromatin re-wetting, and ultimately cell survival, will occur following wound-induced mechanical stress.

### Transient chromatin-lamina dewetting in Re-Wet cells is triggered by rapid tensional relaxation

To identify potential mechanical triggers of transient chromatin-lamina dewetting in Re-Wet cells, we analyzed their spatial distribution after wounding, leveraging the natural tension anisotropy between the anteroposterior (AP) and dorsoventral (DV) axes of the *Drosophila* epithelium (*27*, *28*). Probability maps revealed that Re-Wet cells (TC2) are located farther from the wound than De-Wet cells (TC1), which cluster closer to the laser cut line (**Fig. 2A-B, S3A**). Re-Wet cells extend up to 30 μm from the wound and exhibit an anisotropic distribution aligned with the embryonic DV axis. This is the direction of the largest tissue relaxation following wounding, consistent with pre-existing tension anisotropy in the epithelium. Indeed, the probability distribution of Re-Wet cells (TC2) matches the deformation field generated by the wound in the epidermis, with Re-Wet cells appearing in regions of high deformation and tensional changes (**Fig. S3B**). Dissecting along the DV axis results in elongated wound openings and smaller tissue relaxation due to the mechanical anisotropy (**Fig. 2C**). In these embryos, De-Wet cells remain localized along the cut line, supporting the idea that they result from local damage (**Fig. 2D** and **S3A**). In contrast, Re-Wet cells still distribute along the DV axis, now more symmetrically around the wound: Here as well, the Re-Wet cells distribution matches the deformation field that is more homogeneous around the wound, further supporting a role for rapid tension release in initiating transient chromatin-lamina dewetting (**Fig. 2D, S3A-B**).

**Figure 2:**
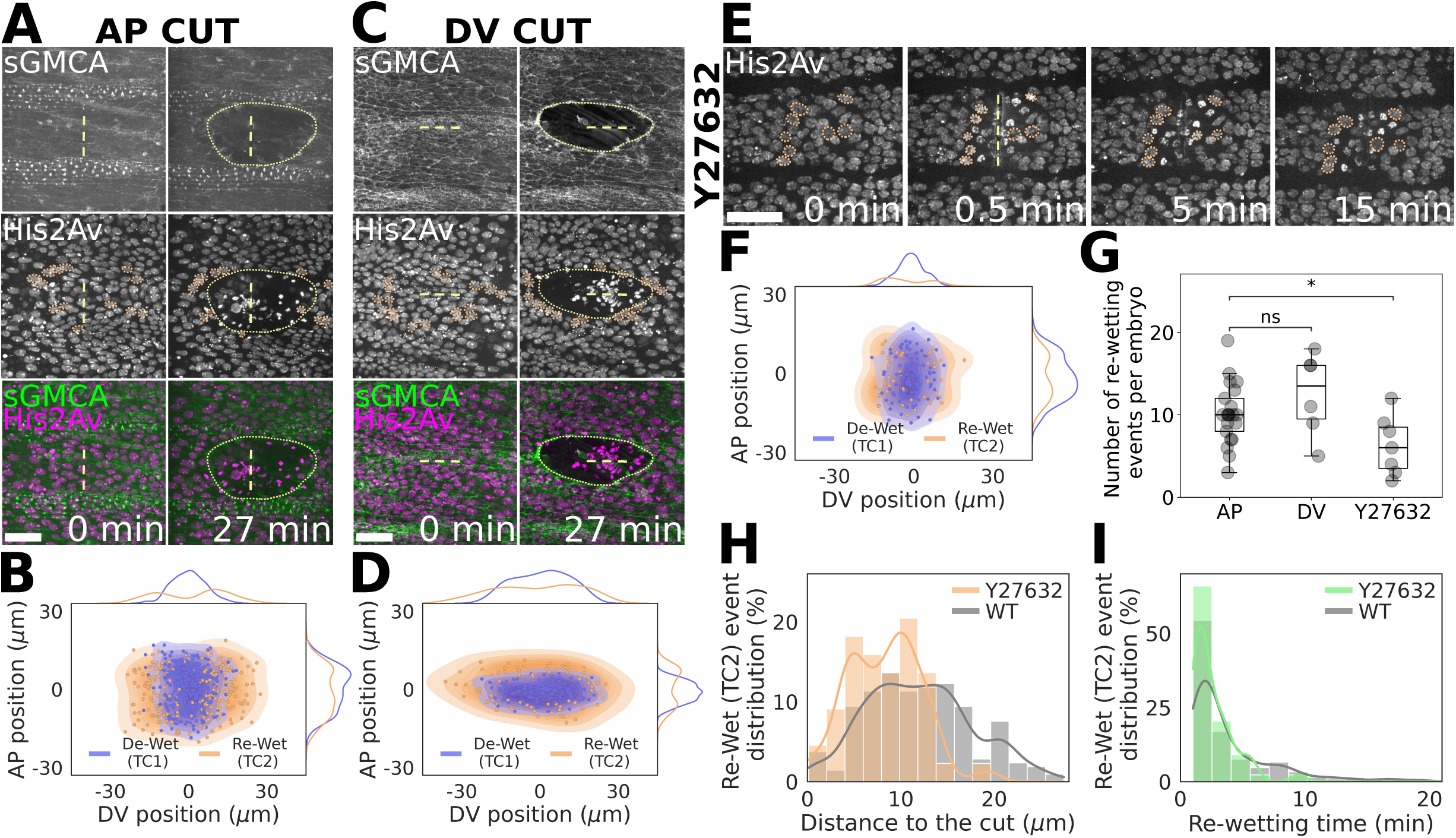
Fast tensional release triggers the reversible chromatin-lamina dewetting. **(A)** Images showing the location of the re-wetting events and the circular wound shape in embryos where the wound was induced along the AP direction, expressing sGMCA and His2Av-mRFP. Scale bar 20 μm. **(B)** Probability distribution map of De-Wet (TC1) and Re-Wet (TC2) cells with respect to the wound induced along the AP direction (*N* = 21 embryos, *n_TC1_* = 278 cells, *n_TC2_* = 212 cells). **(C)** Images showing the location of the re-wetting events and the elliptical wound shape in embryos where the wound was induced along the DV direction, expressing sGMCA and His2Av-mRFP. Scale bar 20 μm. **(D)** Probability distribution map of De-Wet (TC1) and Re-Wet (TC2) cells with respect to the wound induced along the DV direction (*N* = 6 embryos, *n_TC1_* = 88 cells, *n_TC2_* = 75 cells). **(E)** Time-lapse images showing reduced tissue retraction after wounding and closer localization of re-wetting events in Rho-kinase inhibited embryos expressing His2Av-mRFP. Scale bar 20 μm. **(F)** Probability distribution map of De-Wet (TC1) and Re-Wet (TC2) cells with respect to the wound induced in Rho-kinase inhibited embryos (*N* = 8 embryos, *n_TC1_* = 100 cells, *n_TC2_* = 44 cells). **(G)** Occurrence of Re-Wet (TC2) cells in AP-induced cuts, DV-induced cuts and Rho-kinase inhibited embryos. The number of Re-Wet cells is decreased by half in Rho-kinase treated embryos (*N_AP_* = 21 embryos, *N_DV_* = 6 embryos, *N_Y27_* = 8 embryos, p-values: *p_DV_* = 0.51, *p_Y27_ =* 0.0294, one-way ANOVA with Dunnett’s post hoc test versus AP). **(H)** Histogram of Re-Wet (TC2) cell distance to the wound in Rho-kinase inhibited embryos showing the events are localized closer to the wound (*N_WT_* = 21 embryos, *n_WT_* = 212 cells, *N_Y27_* = 8 embryos, *n_Y27_* = 44 cells, *p* < 0.0001, one-tailed *t* test). **(I)** Histogram of re-wetting times of Re-Wet (TC2) cells in control and Rho-kinase inhibited experiments (*N_WT_* = 21 embryos, *n_WT_* = 212 cells, *N_Y27_* = 8 embryos, *n_Y27_* = 44 cells, *p* = 0.027, two-tailed Mann-Whitney U test).

Reducing epithelial tension with the Rho-kinase inhibitor Y-27632 (see Methods) lowered wound recoil and slowed relaxation (**Fig. 2E**, **S3B** and **Movie S4**). Re-Wet cell frequency fell by ∼50%, compared to WT embryos (**Fig. 2G**). They are spatially distributed in regions closer to the wound, with 95% of the Re-Wet cells within 13.3 𝜇𝑚 from the wound compared to 21.8 𝜇𝑚 for WT (**Fig. 2F, H,** and **S3A-B**). However, the timing of re-wetting onset remained similar to WT embryos (**Fig. 2I**). The Y-27632 perturbation experiments therefore indicate that a reduction in contractility and tissue surface tension lessens the occurrence of transient chromatin-lamina dewetting and restricts it in close wound regions, in which some rapid tensional unloading may still occur during wounding.

Altogether, these observations indicate that high tensional relaxation is essential to trigger transient chromatin-lamina dewetting in Re-Wet cells, whereas De-Wet cells likely result from direct damage along the wound cut.

### Chromatin re-wetting is mechanosensitive and requires the LINC complex

To further investigate the mechanical regulation of Re-Wet cells, we applied ectopic mechanical perturbations by compressing embryos, following a previously established protocol (*29*). Compression alters embryo aspect ratio, stretching the epidermis and increasing friction with the vitelline envelope due to tighter contact (**Fig. S3C**). After 20 minutes of compression, we performed AP-axis laser ablations (**Fig. 3A-B**).

**Figure 3:**
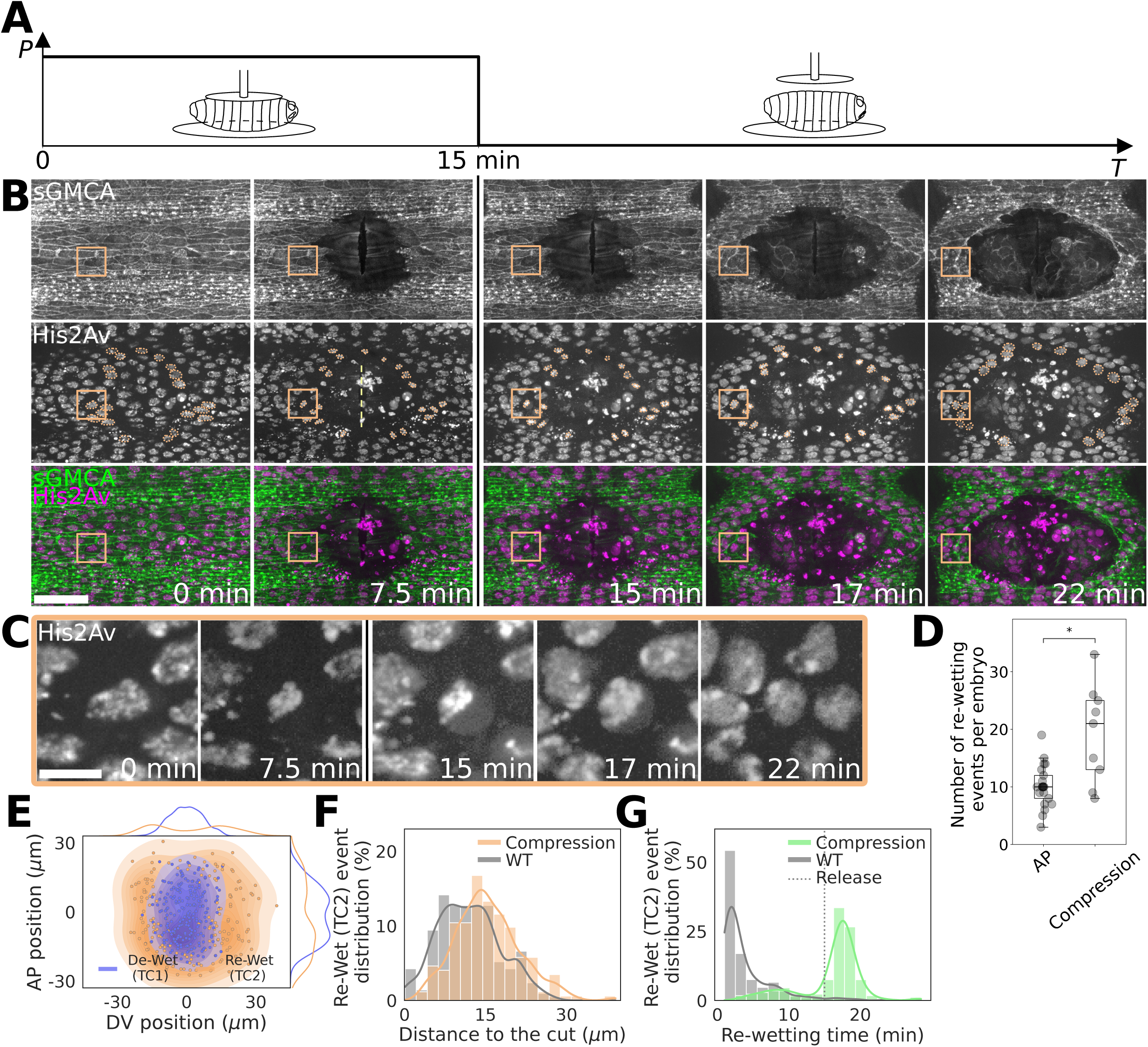
Chromatin re-wetting is mechanosensitive. **(A)** Schematic illustrating experiments conducted with external force application, in which a compression was exerted with a micromanipulator on the embryo for 15 minutes after a cut was induced. **(B)** Time-lapse images showing the suppressed re-wetting while embryos are under compression in embryos expressing sGMCA and His2Av-mRFP. The re-wetting was delayed and almost entirely suppressed until the pressure was released. Re-Wet (TC2) cells are outlined in orange. Scale bars 25 μm. **(C)** Time-lapse images showing a delayed decompaction of a Re-Wet (TC2) cell while the embryo is under compression. Scale bar 5 μm. **(D)** Number of Re-Wet (TC2) events showing a 2x increase in compression experiments compared to the controls. (*N_AP_* = 21 embryos, *N_COMP_ =* 9 embryos*, p_TC2_ =* 0.012, two-tailed *t* test). **(E)** Probability distribution map of De-Wet (TC1) and Re-Wet (TC2) cells with respect to the wound (*N* = 9 embryos, *n_TC1_* = 228 cells, *n_TC2_* = 173 cells). **(F)** Histogram of Re-Wet (TC2) cell distance to the wound in compression experiments (*N_WT_* = 21 embryos, *n_WT_* = 212 cells, *N_COMP_* = 9 embryos, *n_COMP_* = 173 cells, *p* < 0.0001, one-tailed Mann-Whitney U test). **(G)** Histogram of re-wetting times of Re-Wet (TC2) cells in control and compression experiments (*N_WT_* = 21 embryos, *nWT* = 212 cells, *N_COMP_* = 9 embryos, *n_COMP_* = 173 cells, *p* < 0.0001, two-tailed Mann-Whitney U test) showing a suppressed re-wetting until the release of pressure.

Under compression, chromatin-lamina dewetting was still observed around the wound, similarly to uncompressed embryos (**Fig. 3B-C** and **Movie S5**). However, chromatin re-wetting was almost entirely suppressed during compression, even up to 15 minutes of compression, while the average re-wetting time is 4.02 ± 3.8 min in uncompressed embryos (**Fig. 3B-C**). Upon release of compression, as friction dissipated and tissue relaxed, chromatin re-wetting occurred synchronously with tissue relaxation (**Fig. 3B–C** and **G** and **Movie S5)**. Notably, the spatial distribution of Re-Wet cells was broader and more isotropic after compression release (**Fig. 3D-E-F** and **S3A**), confirming that mechanical cues promote their occurrence. These results highlight that changes in the mechanical state of the tissue modulate both the timing and spatial emergence of Re-Wet cells.

To understand how such mechanical cues are transmitted to the chromatin, we tested whether the nuclear–cytoskeletal connection via the LINC complex is required for chromatin re-wetting. We performed wound healing experiments in homozygous Msp300^ΔKASH^ mutants, which lack the KASH domain of the nesprin ortholog Msp300 and therefore disrupt LINC-mediated coupling (*30*). These mutants showed reduced tissue retraction after wounding, consistent with decreased epithelial tension (**Fig. 4A, S3B** and **Movie S6**). Re-Wet cells were still present, but at significantly lower frequency (∼6.6 events/embryo) and located slightly closer to the wound, possibly due to the attenuated tensional changes (**Fig. 4B-C-D** and **S3A-B**). Interestingly, chromatin re-wetting was delayed in Msp300^ΔKASH^ embryos. Many cells remained in the dewetting state for extended periods, with over 30% of re-wetting events occurring only after 10 minutes (compared to only 7% for WT) (**Fig. 4-E**). This indicates that nuclear–cytoskeletal coupling via the LINC complex is required for efficient and timely chromatin re-wetting. It was recently shown that the mechanical coupling between the cytoskeleton, the nuclear envelope, and the chromatin depends on Ataxia Telangiectasia and Rad3-related protein (ATR) (*21*, *31*). To investigate whether ATR is involved in the reversible chromatin-lamina dewetting, we injected embryos with the ATR inhibitor ceralasertib (see Methods). In these embryos, we observed overall a similar number of chromatin-lamina dewetting events (27.2 ± 7.8 compared to 23.3 ± 4.2 on average in control embryos), located slightly closer to the wound (**Fig. 4F-G-I** and **S3A**). However, we observed almost half of the re-wetting events compared to wild type, with many showing delays, indicating that ATR is required for the chromatin re-expansion and wetting of the lamina (**Fig. 4H and J**).

**Figure 4:**
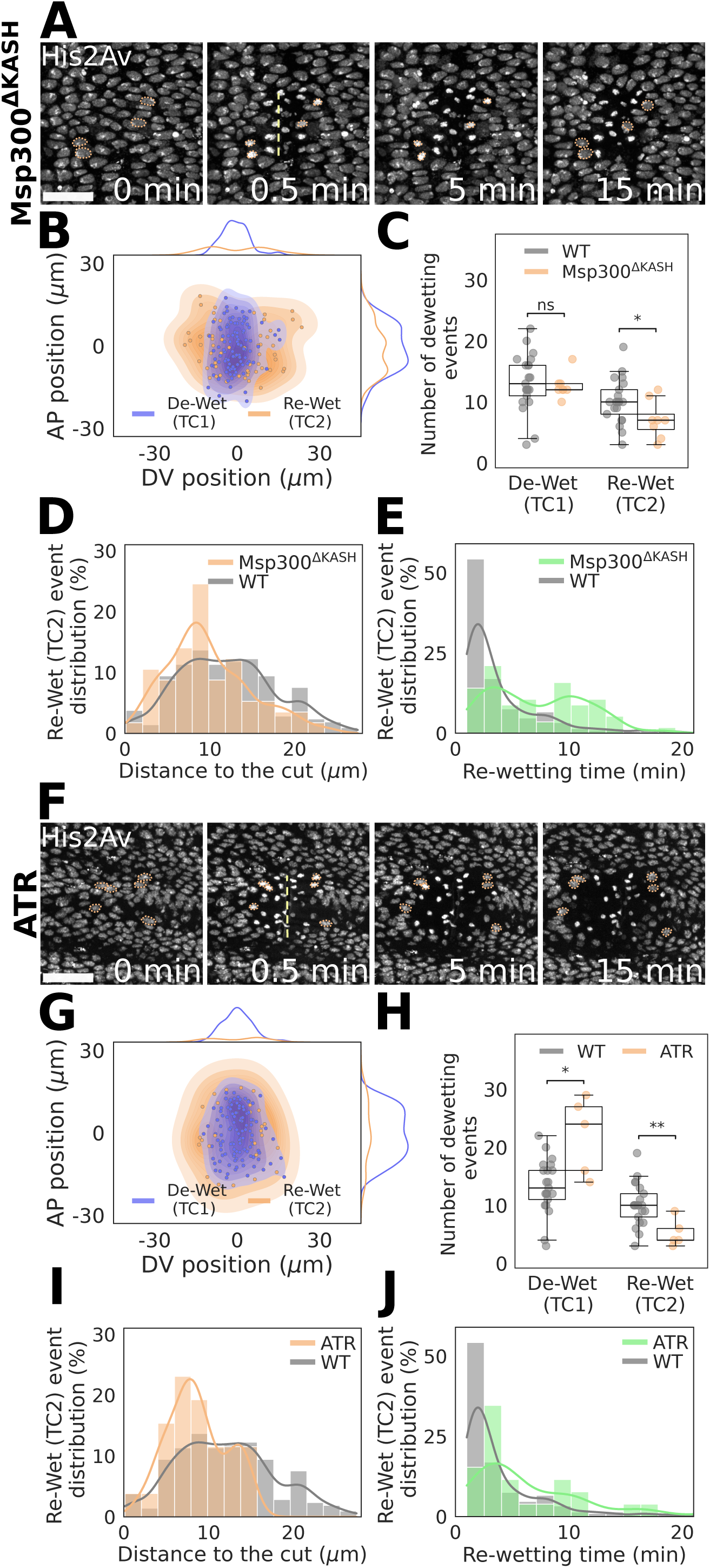
Chromatin re-wetting depends on the LINC complex, and the re-expansion is mediated through ATR. **(A)** Time-lapse images showing reduced tissue retraction after wounding and decreased occurrence of re-wetting events in homozygous Msp300^ΔKASH^ mutants expressing His2Av-mRFP. Scale bar 20 μm. **(B)** Probability distribution map of De-Wet (TC1) and Re-Wet (TC2) cells with respect to the wound (*N* = 8 embryos, *n_TC1_* = 101 cells, *n_TC2_* = 57 cells). **(C)** Number of de-wetting events (TC1 and TC2) showing no change in TC1 occurrence and a decrease in Re-Wet (TC2) occurrence in Msp300^ΔKASH^ mutants compared to the controls. (*N_WT_* = 21 embryos, *N_ΔKASH_* = 8 embryos, p-value*: p_TC1_ =* 0.6254, *p_TC2_ =* 0.0442, two-tailed *t* test) **(D)** Histogram of Re-Wet (TC2) cell distance to the wound in the mutants (*N_WT_* = 21 embryos, *n_WT_* = 212 cells, *N_ΔKASH_* = 8 embryos, *n_ΔKASH_* = 57 cells, *p* = 0.0022, one-tailed *t* test). **(E)** Histogram of re-wetting times of Re-Wet (TC2) cells in control and Msp300^ΔKASH^ mutants experiments (*N_WT_* = 21 embryos, *n_WT_* = 212 cells, *N_ΔKASH_* = 8 embryos, *n_ΔKASH_* = 57 cells, *p* < 0.0001, two-tailed Mann-Whitney U test) showing a delayed re-wetting. **(F)** Time-lapse images showing a decreased occurrence of chromatin-lamina dewetting events, which are localized closer to the wound in ATR-inhibited embryos expressing His2Av-mRFP. Scale bar 20 μm. **(G)** Probability distribution map of De-Wet (TC1) and Re-Wet (TC2) cells with respect to the wound (*N* = 5 embryos, *n_TC1_* = 110 cells, *n_TC2_* = 26 cells). **(H)** Number of dewetting events (TC1 and TC2) showing an increase in TC1 occurrence and a decrease in TC2 occurrence in ATR-inhibited embryos compared to the controls. (*N_WT_* = 21 embryos, *N_ATR_* = 5 embryos*, p_TC1_* = 0.0391, *p_TC2_ =* 0.005, two-tailed *t* test). **(I)** Histogram of TC2 cell distance to the wound in ATR-inhibited embryos (*N_WT_* = 21 embryos, *n_WT_* = 212 cells, *N_ATR_* = 5 embryos, *n_ATR_* = 26 cells, *p* < 0.0001, one-tailed *t* test) showing the cells are located closer to the wound. **(J)** Histogram of re-wetting times of TC2 cells in control and ATR-inhibited embryos showing a slightly delayed re-wetting. (*N_WT_* = 21 embryos, *n_WT_* = 212 cells, *N_ATR_* = 5 embryos, *n_ATR_* = 26 cells, *p* = 0.0002, two-tailed Mann-Whitney U test)

Altogether, these experiments demonstrate that Re-Wet cells are mechanosensitive: a fast release of tissue-scale tension triggers chromatin-lamina dewetting, while subsequent mechanical inputs transmitted through the LINC complex are essential for an ATR-dependent chromatin re-engagement with the lamina.

### Calcium is necessary for chromatin dewetting, stored in the chromatin and released during re-wetting

Wound generation does not only involve a fast mechanical change in the epithelium but also a dramatic change in ionic content in the tissue, particularly calcium (*2–5*). To investigate the impact of calcium on the reversible chromatin-lamina dewetting process, we imaged embryos with genetically encoded calcium sensor Gcamp8, which emit GFP when bound to calcium. As previously reported, the generation of a wound in the epidermis triggers a rapid wave of calcium spreading in the epidermis (**Fig. 5A**). Co-imaging the calcium sensor Gcamp8 and His2Av-mRFP, we could quantify calcium levels within Re-Wet cells (**Fig. 5B**): Just after wounding, calcium levels increase in these cells and then decrease significantly compared to the neighboring cells, indicating a depletion of free intracellular calcium (**Fig. 5B-C** and **S4A** and **Movie S7**). Interestingly, the re-expansion of the chromatin and re-wetting of the lamina correlate with an intranuclear burst of calcium (**Fig. 5B-D** and **Movie S7**), indicating that calcium was stored in the nucleus and chromatin region.

**Figure 5:**
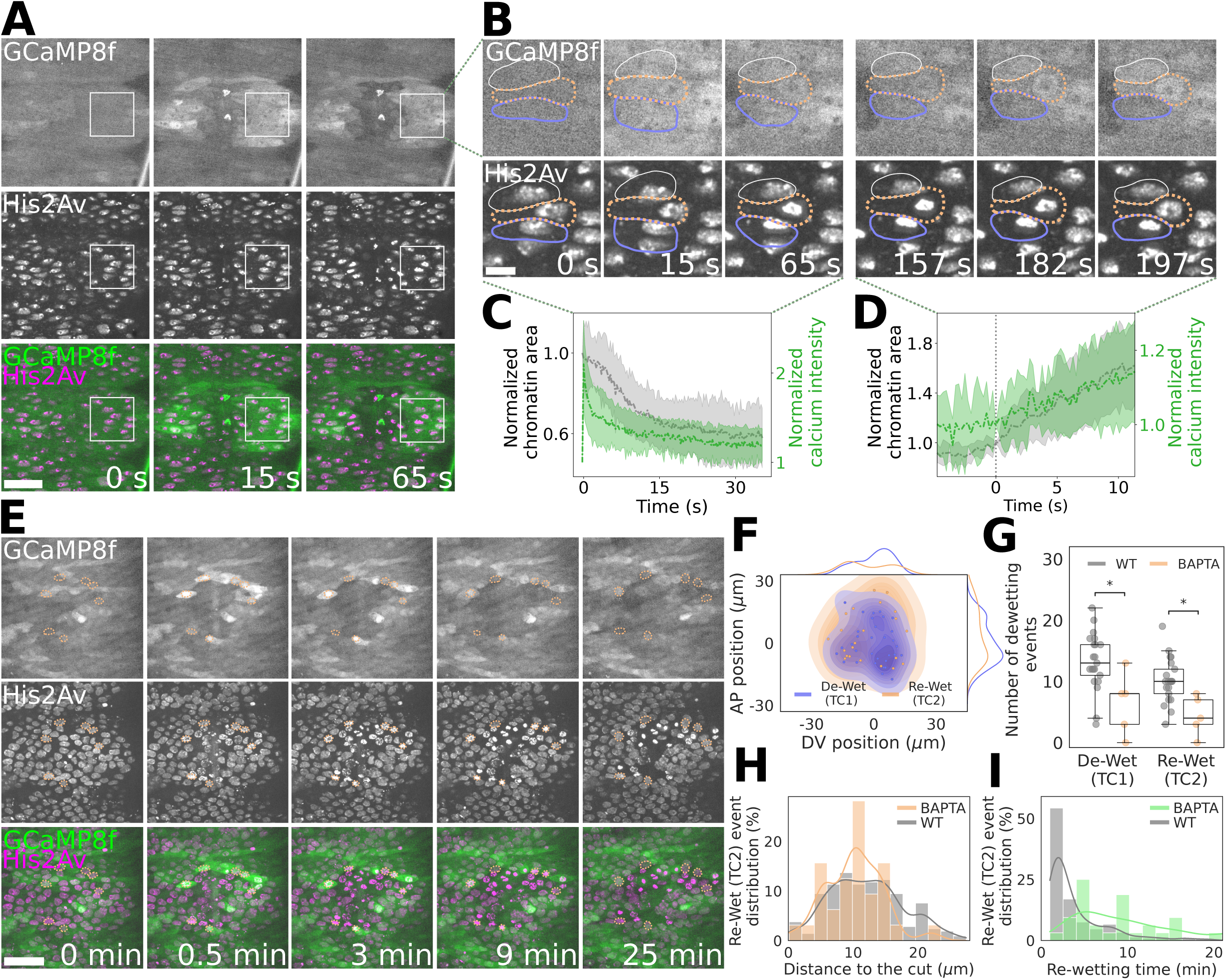
Calcium is necessary for chromatin dewetting, calcium is stored in the chromatin and released during re-expansion. **(A)** Time-lapse images showing a rapid wave of calcium after wound induction in the epidermis of embryos expressing His2Av-mRFP and calcium sensor Gcamp8. Scale bar 20 μm. **(B)** Close-up showing a depletion of free intracellular calcium in Re-Wet (TC2) cells while the chromatin is compacted, after the calcium wave has decayed, subsequently, the chromatin re-expansion correlates with a calcium burst in the nucleus. A De-Wet (TC1) (blue) and an unaffected (white) cell are shown outlined for comparison. Scale bar 5 μm. **(C)** Graph showing normalized chromatin area and normalized calcium intensity in Re-Wet (TC2) cells during the dewetting phase (*N* = 2 embryos, *n* = 10 cells). The following behavior can be observed: rapid increase in calcium intensity during the calcium wave, and a subsequent depletion during the dewetting. The origin of time is set just after cut after the calcium signal stabilized. **(D)** Normalized chromatin area and normalized calcium intensity in Re-Wet (TC2) cells during the re-wetting phase (*N* = 3 embryos, *n* = 10 cells). Graph showing concurrent increase in area and the calcium intensity during the re-wetting. The origin of time is set at the initiation of the re-wetting phase. **(E)** Time-lapse images showing a reduction of transient lamina-chromatin compaction events in embryos co-expressing His2Av-mRFP and Gcamp8 after chelation of calcium with BAPTA. Scale bar 20 μm. **(F)** Probability distribution map of De-Wet (TC1) and Re-Wet (TC2) cells with respect to the wound in BAPTA treated embryos (*N* = 5 embryos, *n_TC1_* = 32 cells, *n_TC2_* = 22 cells). **(G)** Number of dewetting events (TC1 and TC2) showing a decrease in TC1 and TC2 occurrences in BAPTA treated embryos compared to the controls. (*N_WT_* = 21 embryos, *N_BAPTA_* = 5 embryos*, p_TC1_ =* 0.0339, *pTC2 =* 0.0111, two-tailed *t* test). **(H)** Histogram of TC2 cell distance to the wound in BAPTA treated embryos. (*N_WT_* = 21 embryos, *n_WT_* = 212 cells, *N_BAPTA_* = 5 embryos, *n_BAPTA_* = 22 cells, *p* = 0.3989, two-tailed *t* test) **(I)** Histogram of re-wetting times of TC2 cells in control and BAPTA treated embryos showing a slightly delayed re-wetting (*NWT* = 21 embryos, *n_WT_* = 212 cells, *NBAPTA* = 5 embryos, *n_BAPTA_* = 22 cells, *p* < 0.0001, two-tailed Mann Whitney U test).

We then decided to deplete free calcium in the epithelium using the cell-permeant calcium chelator BAPTA AM (see Methods). Upon chelation of calcium, wound generation shows a strong reduction in events of chromatin compaction and of reversible chromatin-lamina dewetting (10.8 ± 7.5 of total compaction events compared to 23.3 ± 4.2 in WT) (**Fig. 5E-G** and **Movie S8**). Instead, in 50% of the affected cells, the chromatin partially compacts and remains associated with the lamina with no apparent dewetting (**Fig. 5E** and **Fig. S4C-D**). The few events of reversible chromatin-lamina dewetting similar to WT observed after wounding are still associated with an increase in free calcium levels remaining from BAPTA treatment (**Fig. 5E** and **S4B**), indicating that calcium is necessary for the compaction to occur. Moreover, the distribution of De-Wet and Re-Wet cells now overlaps **(Fig. 5F** and **S3A**), and, while the spatial distribution of the Re-Wet cells appears unaffected (**Fig. 5H**), the timing of chromatin re-expansion is strongly affected with 10-20 minutes time delays (average re-expansion time of 9.07 ± 5.6 min) (**Fig. 5I**).

Overall, our results indicate that calcium is essential for chromatin compaction and chromatin lamina dewetting after wound generation. It indicates that calcium is stored in the nucleus when the chromatin compacts and is released at the moment of decompaction in a transient burst of calcium.

### Chromatin-lamina re-wetting triggers an actomyosin contraction and promotes activation of the JnK pathway and restoration of tensile healing forces

Do the reversible chromatin-lamina dewetting events impact the mechanics of tissues and the healing response of the epithelia? To address this question, we imaged the chromatin (His2Av-mRFP) and the actomyosin cytoskeleton (sGMCA) simultaneously: As a response to wounding, the actomyosin reorganizes in a supracellular cable surrounding the wound that will power the healing response. By imaging the actin and myosin in Re-Wet cells, we could observe that the nuclear re-expansion of the chromatin occurs systematically before the onset of the healing response and the formation of the actin cable (**Fig. 6A** and **Movie S9**). Interestingly, by looking closely at the actomyosin cytoskeleton and cellular shape of the cells undergoing chromatin-lamina re-wetting, we could identify a systematic accumulation of actin and myosin at the apical site of the cell and a consequent fast reduction of the apical site following the nuclear lamina re-wetting (**Fig. 6A-B** and **Fig S4F**). It is known that a sudden intracellular calcium increase triggers an actomyosin contraction (*32*). The calcium burst observed during the chromatin-lamina re-wetting is therefore likely to be at the origin of the observed consequent actomyosin contraction. This actomyosin contraction always precedes the formation of the supracellular actomyosin cable. After wounding, we can observe an irregular wound-edge due to a reduced line tension for several minutes (**Fig. 6C**). When the chromatin re-expands in the Re-Wet cells, followed by an actomyosin contraction in these cells, a systematic straightening of the leading edge happens in the vicinity of these cells associated with an increase in actin levels at the leading edge, indicating the formation of the actin cable (**Fig. 6C-D**).

**Figure 6:**
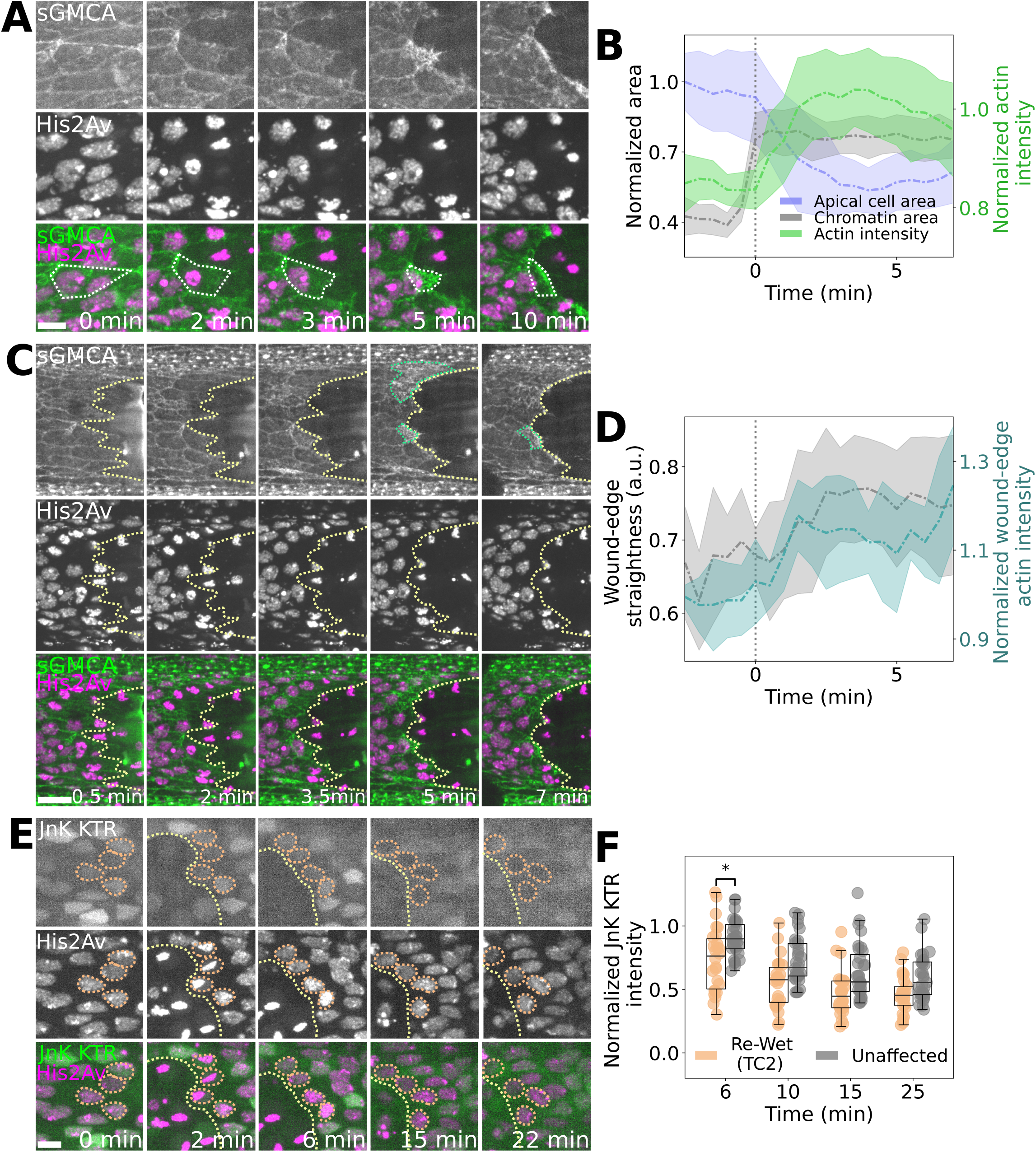
Chromatin re-wetting triggers an actomyosin contraction and promotes the activation of the JnK pathway. **(A)** Time-lapse images showing the accumulation of actin at the apical site of the cell and a fast reduction of the apical site area in an embryo expressing His2Av-mRFP and sGMCA. The white dashed line depicts the apical site of the cells. Scale bar 5 μm. **(B)** Graph showing the normalized chromatin (gray) and apical site (blue) area in Re-Wet (TC2) cells, and the average normalized actin intensity of the Re-Wet (TC2) cells (green) (*N* = 5 embryos, *n* = 39 cells, median + mad). The origin of time is set at the re-wetting time. The accumulation of actin at the apical site of the cell and a fast reduction of the apical site occur after re-wetting. The dashed gray vertical line represents the median re-wetting time. **(C)** Time-lapse images showing a straightening of the leading edge and an increase in actin levels at the leading edge following the actomyosin contraction in an embryo expressing His2Av-mRFP and sGMCA. The yellow dashed line depicts the wound edge, and the cyan dashed line depicts areas with increased actin intensity. Scale bar 10 μm. **(D)** Graph showing the straightening of the leading edge and an increase in actin levels at the leading edge close to Re-Wet (TC2) cells (*N =* 5 embryos, *n =* 24 cells, median + mad). The origin of time is set at the re-wetting time. The dashed gray vertical line represents the re-wetting event. **(E)** Representative images showing an earlier decay in JnK levels in the Re-Wet (TC2) cells (shown in orange dashed outlines), compared to the remaining wound edge cells, which will decay with a time delay. Scale bar 5 μm. **(F)** Boxplots of the normalized JnK intensity in Re-Wet (TC2) and unaffected wound-edge cell nuclei (*N* = 3 embryos, *n_TC2_* = 26 cells, *n_unaffected_* = 31 cells, median shown, two-way ANOVA with Šidák’s multiple comparisons test: factor cell type, *p* = 0.0006; factor time, *p* < 0.0001; interaction, *p =* 0.6927).

This observation suggests that the chromatin-lamina re-wetting and consequent actomyosin contraction promote activation of the healing response in the tissue. Previous in vitro studies reported that nuclear deformations could trigger activation of the Jun Kinase (JnK) pathway (*33*, *34*). In our case, the fast dewetting and re-wetting of the chromatin in the Re-Wet cells is also associated with large nuclear deformation (See **Fig. 1G-H-I**). Possibly, the presence of dramatic changes in chromatin and nuclear shape could trigger the activation of the JnK pathway here as well. JnK is known to be activated during wound healing and to be essential for the healing response (*1*). In order to investigate the activation of the JnK pathway, we used embryos expressing a Jnk activity reporter in the epidermis (JnK KTR) (*35*, *36*). When the JnK is inactive, the reporter is located in the nucleus, and it translocates to the cytoplasm once JnK is activated. We simultaneously monitored the chromatin with His2Av-mRFP and JnK activity with our reporter. We could observe that JnK remains inactive in the entire tissue soon after wounding, when the chromatin compacts in cells neighboring the wound (**Fig. S4E**). To identify potential differences in JnK activation between cells that undergo reversible chromatin lamina dewetting, and other wound edge cells, we quantified JnK KTR in the nucleus after chromatin re-expansion had occurred (**Fig. 6E-F**). We could observe an earlier decay in JnK levels in Re-Wet cells, indicating an early JnK activation in these cells compared to the remaining wound edge cells that will reach similar JnK levels only a few minutes after (**Fig. 6F**).

Altogether, our results indicate that Re-Wet cells are triggering an actomyosin contraction, promoting wound edge straightening, actin ring formation, and JnK activation necessary for tissue repair.

## Discussion

Our study demonstrates that the rapid mechanical unloading that occurs during wound generation triggers a fast and reversible detachment of chromatin from the nuclear lamina, a process we define as chromatin dewetting. This detachment is initiated by a loss of cytoskeletal force transmission through the LINC complex, likely due to actin disassembly, and is facilitated by a concomitant rise in intracellular calcium. Re-wetting requires the reestablishment of tension through the LINC complex, which leads to the release of calcium previously stored in compacted chromatin and subsequently triggers cellular contraction and the healing response. Consistent with previous studies, these findings reveal that chromatin–lamina interactions are not passive structural elements but dynamic, force-sensitive regulators of nuclear mechanics that can rapidly remodel in response to mechanical cues and ultimately influence cellular and tissue mechanical behavior.

Reversible chromatin-lamina dewetting emerges as a previously unrecognized and rapid mode of nuclear mechanosensitivity. This mechanosensing process depends on calcium, which plays a dual role: during dewetting, it accumulates within the chromatin, promoting its compaction and dewetting from the lamina, potentially through direct binding (*37*, *38*) or by activating chromatin-compaction machinery such as condensins or cohesins (*39*, *40*). During re-wetting, calcium is released, triggering actomyosin contraction and contributing to wound edge contraction. ATR is also required for efficient chromatin reattachment, suggesting that it integrates mechanical cues at the chromatin–lamina interface with the structural recovery of the nucleus (*21*, *22*). Functionally, this transient chromatin–lamina uncoupling could serve at least three purposes: (i) mechanical protection of the genome through compaction; (ii) buffering of abrupt force changes at the nuclear envelope; and (iii) activation of downstream cellular responses, including contraction and signaling pathway initiation. This chromatin-lamina dewetting process involves major remodeling of the 3D architecture of the chromatin within a few minutes. It would be interesting to investigate in the future whether these mechanically induced changes in chromatin architecture impact epigenetic regulation and transcriptional programs in these cells, particularly in the context of tissue regeneration.

This mechanosensing mechanism is embedded in a broader multiscale feedback loop that links tissue-level injury to nuclear signaling and chromatin architecture modulation and back again to epithelial remodeling. Injury-induced relaxation of epithelial tension initiates chromatin dewetting and calcium sequestration, followed by re-wetting, calcium release, actomyosin contraction, and JnK pathway activation. These events unfold within minutes of injury and are tightly coordinated, highlighting the responsiveness of the nuclear compartment to mechanical inputs. This is reminiscent of concepts of cellular tensegrity, in which subcellular structures act as tensile-loaded elements that ensure a global mechanical stability at the cellular and tissue level (*41*). Here, the tensegrity elements encompass the cytoskeleton, the nuclear envelope and the chromatin itself. An external perturbation at the tissue scale transiently disrupts the mechanical connectivity between these subcellular elements, which then actively respond by generating forces that propagate through the tissue to restore tensile equilibrium.

Although described in the context of epithelial wound repair, this mechanism is a fundamental cellular mechanism that likely operates, to varying extents, in other biological processes involving rapid mechanical changes, such as spinal cord compression or traumatic brain injury (*42–44*). In such cases, chromatin–lamina dynamics may contribute to nuclear integrity maintenance and early activation of repair pathways. By revealing chromatin dewetting and re-wetting as a rapid and regulated nuclear response, our findings could potentially open new, general protective strategies against acute mechanical stress.

## Materials and methods

### Drosophila Stock

The following Drosophila strains were used: sGMCA, HisRFP (Bloomington stock centre #59023). His2Av-mRFP (Bloomington stock centre #23650), UAS GCamp8f (Bloomington stock centre #92588), Nullo Gal4 (Bloomington stock centre #26875), Lamin::GFP (Bloomington stock centre #6837), Msp300^ΔKash^ (Bloomington stock centre #26781), UAS JnK GFP (reference Amelie’s paper).

### Embryo collection and sample preparation

Embryos were collected at 22 or 25°C and aged overnight. Following collection, embryos were dechorionated in 5% bleach for two minutes and subsequently thoroughly washed with water for one minute. The embryos were mounted ventrally on a coverslip with heptane glue for conventional confocal microscopy (ventral side of the embryo was facing the glue) and covered with halo carbonated oil to prevent them from drying.

### Imaging

Imaging of the samples was performed on an Olympus IXplore SpinSR10 spinning disk confocal super-resolution microscope equipped with a 100x/1.5 NA oil immersion objective. Fluorescence images were acquired with the excitation confocal laser lines of 488 nm, 100 mW, Diode, and 561 nm, 100 mW, Diode. Time-lapse z-stack images were acquired at 30 s time intervals with a step-size of 1 µm; except for faster time resolution used to image the Lamin-GFP, with the time intervals being 10 s. For calcium acquisition, a single z-slice was imaged every 252 ms. All experiments were conducted at 25 °C (or at 22 °C for the imaging in Fig. 1I and Fig. 2A-C) with 15% laser power for both 488 nm and 561 nm.

### Laser dissection

Laser dissection was carried out using the Olympus IXplore SpinSR10, equipped with a Rapp OptoElectronic UGA-42 Caliburn 355/42 pulsed laser (355nm, 1kHz, 42µJ/pulse) with a UGA-42 Geo module. Cuts were performed at 10 – 20 % of the laser power, dissecting a 25 µm straight line, with repetition of dissection of 10 – 30 times. All cuts have been induced along the AP axis, unless stated otherwise, to compare all perturbation experiments against the controls. The cuts were located between two segment boundaries on the ventral side of the embryo in the late embryonic stage.

### External force application

A compression device consisting of a metal weight on a coverslip, weighing 0.92 ± 0.01 g, was used to apply an external force on the embryos. Two stripes of tape (100 µm thick) served as spacers to ensure uniform and reproducible compression from embryo to embryo. Compression was initiated 20 minutes before the acquisition to ensure a stable compression on the embryo. The compression was maintained for 15 minutes after laser dissection, and the pressure was manually released thereafter.

### Microinjection

Embryos were dehydrated for 10 minutes before covering them with halo carbonated oil and then mounted on a coverslip covered with heptane glue. Injection of pharmaceutical drugs was performed in the vitelline space from the posterior side of the embryo. The following pharmaceutical inhibitors were used: 5 mM Rho-kinase inhibitor (Y27632, Sigma Aldrich SCM075), 2 mM extracellular Ca2+ chelator BAPTA, and 1mM ATR inhibitor ceralasertib.

### Mechanical wound generation experiment

Embryos were punctured with a micropipette held on a binocular microscope to induce mechanically generated wounds. The micropipette was rapidly inserted deep in the embryo and removed to ensure epidermal damage. The embryos were then imaged on the Olympus IXplore SpinSR10 spinning disk.

### Image analysis

Time-lapse images were analyzed using Fiji (ImageJ) and Python. Unless otherwise stated, analyses were performed on maximum-intensity projected z-stacks.

Nuclei were segmented and stitched with Cellpose (*45*), tracked over time with TrackMate when required (*46*), and manually corrected. Labeled nuclei were classified into three groups: De-Wet (TC1), Re-Wet (TC2), and unaffected cells. Cell classification was based on the temporal gradients of chromatin area and mean fluorescence intensity within each mask. De-Wet (TC1) cells exhibited a single rapid peak in the gradients but did not show a second pronounced peak, whereas Re-Wet (TC2) cells displayed multiple prominent peaks. Unaffected cells showed no prominent peaks in either the area or intensity gradients over time.

The position of the laser cut was manually determined from the ablation imprint on the maximum projection. Chromatin positions were calculated as centroids of the chromatin masks and subsequently rotated relative to the cut to ensure alignment across embryos.

Re-wetting times of Re-Wet (TC2) cells were estimated from the temporal profile of chromatin area. For each cell, the re-wetting time was defined as the time point corresponding to the peak in the gradient (first derivative) of the area curve, reflecting the moment of maximal change of area.

The chromatin–wound distances were measured as the distance between the centroid of the chromatin mask at the first frame (*t* = 0 min, before the dissection was performed) and the laser cut. As the ablation was not a point cut, distances were computed as follows: If the perpendicular projection of the centroid position onto the laser cut line fell outside the endpoints, the distance to the nearest endpoint was used; otherwise, the perpendicular distance to the line was taken.

Histograms of chromatin-wound distances and re-wetting times were generated with bin sizes of 2 μm and 1.5 min, respectively. Kernel density estimation maps of De-Wet (TC1) and Re-Wet (TC2) cell positions relative to the wound were generated to visualize spatial distributions. Kernel density estimation was performed using a Gaussian kernel with bandwidth automatically determined by Scott’s rule.

Positions of nuclei before and after the cut were used to compute the deformation field around the wound. Displacement magnitudes were then mapped onto a downsampled grid and smoothed with a Gaussian filter to generate a heatmap, in which intensity reflects local displacement.

To analyze chromatin and nuclear envelope dynamics, lamin and histone signals were segmented and stitched with Cellpose in 3D over time. Nuclear area and circularity were measured from the midplane section (defined as the plane of maximal nuclear/chromatin area). Circularity was calculated as 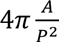, where 𝐴 is area and 𝑃 is perimeter.

For cytoplasmic GFP and NLS-GFP, cells and nuclei, respectively, were manually segmented. Fluorescence signals were corrected for photobleaching by exponential fitting, background-subtracted (background estimated from non-embryonic regions), and quantified as the mean intensity per object.

Calcium intensity was calculated as the mean fluorescence intensity of Gcamp8 expressing embryos within the chromatin mask of Re-Wet (TC2) and unaffected cells. To compare across different cells with different re-wetting times, the reference time was defined as the initiation of re-wetting (initiation of the increase in chromatin area).

To assess whether calcium is required for chromatin compaction, calcium levels were analyzed in BAPTA-injected embryos. Chromatin of Re-Wet (TC2) and unaffected cells was segmented as described above. Fluorescence signals were corrected for photobleaching by exponential fitting. The mean calcium intensity within each chromatin mask was measured at the first time point before cuts and second time point just after cut, and the increase was calculated as the difference between these values.

To analyze the actomyosin ring formation, the measurement of intensity and area of chromatin, and the apical site of the cell were performed on the time-lapse images of His2Av-mRFP and sGMCA expressing embryos. Chromatin and apical cell sites of Re-Wet (TC2) and unaffected cells were segmented over time manually. Fluorescence intensity was quantified as the mean signal within each mask. To compare Re-Wet (TC2) cells with different re-wetting times, measurements of Re-Wet (TC2) cells were aligned to a reference time defined as the conclusion of re-wetting.

The leading edge of the wound was manually segmented over time to analyze its dynamics. For each Re-Wet (TC2) cell, 100-pixel segments of the leading edge were extracted at each time point. Two parameters were quantified: (i) straightness of the segment and (ii) mean actin intensity over a dilated leading-edge mask. Straightness was defined as 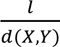, where 𝑙 is the length of the segment, and 𝑑(𝑋, 𝑌) is the maximal Euclidean distance between any two points of the segment. To compare across different cells with different re-wetting times, the reference time was defined as the moment of re-wetting occurrence.

To analyze whether the JnK pathway is activated earlier in the Re-Wet (TC2) cells compared to the unaffected ones, the nucleus was segmented manually. The intensity was corrected for photobleaching by correcting against the mean signal in unaffected cells far from the wounded epidermis. The mean JnK KTR intensity within each nuclear mask was measured at different time points.

### Statistical analysis

Normality was assessed using the D’Agostino-Pearson or the Shapiro–Wilk test, based on sample size. Normally distributed datasets were analyzed with unpaired Student’s *t*-tests with Welch’s correction, while non-normally distributed datasets were analyzed with Mann–Whitney tests. Depending on the hypothesis, one- or two-tailed tests were applied as indicated. Comparisons among three groups were performed using one-way ANOVA with Dunnett’s post hoc test to evaluate differences relative to the control group, and two-way repeated-measures ANOVA was used for experiments involving two independent variables, followed by post hoc multiple comparisons. Significance: ns = not significant, \**p*<0.05, \*\**p*<0.01, \*\*\**p*<0.001 and \*\*\*\**p*<0.0001.

## Acknowledgements.

We thank Manuel Mendoza, Alberto Elosegi-Artoga, Nimesh Chahare, Juan Francisco Abenza Martínez, Amy Beedle, Miguel González Martín, and Pere Roca-Cusachs for critical reading of the manuscript, and the BRALM microscopy core facility. We thank Prof. Marco Foiani for insightful discussions on ATR and ATM. The research leading to these results has received funding from the Spanish Ministry of Science and Innovation, Plan National, PID2019-109117GB-100, PID2022-142779NB-100 and PID2022-141040NA-I00 and from the Basque government, Proyectos de Investigacion Basica/Aplicada, PIBA_2023_1_0033. This work was supported in part by the Fundación Biofisika Bizkaia.

## References

1. G. C. Gurtner, S. Werner, Y. Barrandon, M. T. Longaker, Wound repair and regeneration. Nature 2008 453:7193 453, 314–321 (2008).

2. S. Restrepo, K. Basler, Drosophila wing imaginal discs respond to mechanical injury via slow InsP3R-mediated intercellular calcium waves. Nature Communications 2016 7:1 7, 1–9 (2016).

3. E. K. Shannon, A. Stevens, W. Edrington, Y. Zhao, A. K. Jayasinghe, A. Page-McCaw, M. S. Hutson, Multiple Mechanisms Drive Calcium Signal Dynamics around Laser-Induced Epithelial Wounds. Biophys J 113, 1623 (2017).

4. S. K. Yoo, C. M. Freisinger, D. C. LeBert, A. Huttenlocher, Early redox, Src family kinase, and calcium signaling integrate wound responses and tissue regeneration in zebrafish. J Cell Biol 199, 225–234 (2012).

5. W. Razzell, I. R. Evans, P. Martin, W. Wood, Calcium Flashes Orchestrate the Wound Inflammatory Response through DUOX Activation and Hydrogen Peroxide Release. Current Biology 23, 424 (2013).

6. W. Wood, A. Jacinto, R. Grose, S. Woolner, J. Gale, C. Wilson, P. Martin, Wound healing recapitulates morphogenesis in Drosophila embryos. Nat Cell Biol 4, 907–912 (2002).

7. M. Antunes, T. Pereira, J. V. Cordeiro, L. Almeida, A. Jacinto, Coordinated waves of actomyosin flow and apical cell constriction immediately after wounding. Journal of Cell Biology 202, 365–379 (2013).

8. P. Martin, J. Lewis, Actin cables and epidermal movement in embryonic wound healing. Nature 360, 179–183 (1992).

9. M. Rämet, R. Lanot, D. Zachary, P. Manfruelli, JNK signaling pathway is required for efficient wound healing in Drosophila. Dev Biol 241, 145–156 (2002).

10. M. Bosch, F. Serras, E. Martín-Blanco, J. Baguñà, JNK signaling pathway required for wound healing in regenerating Drosophila wing imaginal discs. Dev Biol 280, 73–86 (2005).

11. G. Nikoloudaki, S. Brooks, A. P. Peidl, D. Tinney, D. W. Hamilton, JNK Signaling as a Key Modulator of Soft Connective Tissue Physiology, Pathology, and Healing. Int J Mol Sci 21, 1015 (2020).

12. M. J. Lee, M. R. Byun, M. Furutani-Seiki, J. H. Hong, H. S. Jung, YAP and TAZ regulate skin wound healing. J Invest Dermatol 134, 518–525 (2014).

13. L. Zechini, C. Amato, A. Scopelliti, W. Wood, Piezo acts as a molecular brake on wound closure to ensure effective inflammation and maintenance of epithelial integrity. Curr Biol 32, 3584–3592.e4 (2022).

14. B. Enyedi, M. Jelcic, P. Niethammer, The Cell Nucleus Serves as a Mechanotransducer of Tissue Damage-Induced Inflammation. Cell 165, 1160–1170 (2016).

15. S. Mascharak, M. Griffin, H. E. Talbott, J. L. Guo, J. Parker, A. G. Morgan, C. Valencia, M. M. Kuhnert, D. J. Li, N. E. Liang, R. M. Kratofil, J. A. Daccache, I. Sidhu, M. F. Davitt, N. Guardino, J. M. Lu, D. B. Abbas, N. M. D. Deleon, C. V. Lavin, S. Adem, A. Khan, K. Chen, D. Henn, A. Spielman, A. Cotterell, D. Akras, M. D. Mauricio Downer, R. Tevlin, H. P. Lorenz, G. C. Gurtner, M. Januszyk, S. Naik, D. C. Wan, M. T. Longaker, Inhibiting mechanotransduction prevents scarring and yields regeneration in a large animal model. Sci Transl Med 17 (2025).

16. S. Dupont, L. Morsut, M. Aragona, E. Enzo, S. Giulitti, M. Cordenonsi, F. Zanconato, J. Le Digabel, M. Forcato, S. Bicciato, N. Elvassore, S. Piccolo, Role of YAP/TAZ in mechanotransduction. Nature 2011 474:7350 474, 179–183 (2011).

17. A. Elosegui-Artola, I. Andreu, A. E. M. Beedle, A. Lezamiz, M. Uroz, A. J. Kosmalska, R. Oria, J. Z. Kechagia, P. Rico-Lastres, A. L. Le Roux, C. M. Shanahan, X. Trepat, D. Navajas, S. Garcia-Manyes, P. Roca-Cusachs, Force Triggers YAP Nuclear Entry by Regulating Transport across Nuclear Pores. Cell 171, 1397–1410.e14 (2017).

18. A. J. Lomakin, C. J. Cattin, D. Cuvelier, Z. Alraies, M. Molina, G. P. F. Nader, N. Srivastava, P. J. Saez, J. M. Garcia-Arcos, I. Y. Zhitnyak, A. Bhargava, M. K. Driscoll, E. S. Welf, R. Fiolka, R. J. Petrie, N. S. de Silva, J. M. González-Granado, N. Manel, A. M. Lennon-Duménil, D. J. Müller, M. Piel, The nucleus acts as a ruler tailoring cell responses to spatial constraints. Science (1979) 370 (2020).

19. V. Venturini, F. Pezzano, F. C. Castro, H. M. Häkkinen, S. Jiménez-Delgado, M. Colomer-Rosell, M. Marro, Q. Tolosa-Ramon, S. Paz-López, M. A. Valverde, J. Weghuber, P. Loza-Alvarez, M. Krieg, S. Wieser, V. Ruprecht, The nucleus measures shape changes for cellular proprioception to control dynamic cell behavior. Science (1979) 370 (2020).

20. M. M. Nava, Y. A. Miroshnikova, L. C. Biggs, D. B. Whitefield, F. Metge, J. Boucas, H. Vihinen, E. Jokitalo, X. Li, J. M. García Arcos, B. Hoffmann, R. Merkel, C. M. Niessen, K. N. Dahl, S. A. Wickström, Heterochromatin-Driven Nuclear Softening Protects the Genome against Mechanical Stress-Induced Damage. Cell 181, 800–817.e22 (2020).

21. A. Kumar, M. Mazzanti, M. Mistrik, M. Kosar, G. V. Beznoussenko, A. A. Mironov, M. Garrè, D. Parazzoli, G. V. Shivashankar, G. Scita, J. Bartek, M. Foiani, ATR mediates a checkpoint at the nuclear envelope in response to mechanical stress. Cell 158, 633–646 (2014).

22. G. Bastianello, G. Porcella, G. V. Beznoussenko, G. Kidiyoor, F. Ascione, Q. Li, A. Cattaneo, V. Matafora, A. Disanza, M. Quarto, A. A. Mironov, A. Oldani, S. Barozzi, A. Bachi, V. Costanzo, G. Scita, M. Foiani, Cell stretching activates an ATM mechano-transduction pathway that remodels cytoskeleton and chromatin. Cell Rep 42 (2023).

23. T. Bhatt, R. Dey, A. Hegde, A. A. Ketkar, A. J. Pulianmackal, A. P. Deb, S. Rampalli, C. Jamora, Initiation of wound healing is regulated by the convergence of mechanical and epigenetic cues. PLoS Biol 20, e3001777 (2022).

24. E. Jokl, A. F. Mullan, K. Simpson, L. Birchall, L. Pearmain, K. Martin, J. Pritchett, S. Raza, R. Shah, N. W. Hodson, C. J. Williams, E. Camacho, L. Zeef, I. Donaldson, V. S. Athwal, N. A. Hanley, K. Piper Hanley, PAK1- dependent mechanotransduction enables myofibroblast nuclear adaptation and chromatin organization during fibrosis. Cell Rep 42 (2023).

25. S. Hu, D. J. Chapski, N. D. Gehred, T. H. Kimball, T. Gromova, A. Flores, A. C. Rowat, J. Chen, R. R. S. Packard, E. Olszewski, J. Davis, C. D. Rau, T. A. McKinsey, M. Rosa-Garrido, T. M. Vondriska, Histone H1.0 couples cellular mechanical behaviors to chromatin structure. Nature Cardiovascular Research 2024 3:4 3, 441–459 (2024).

26. G. Reiter, Dewetting of thin polymer films. Phys Rev Lett 68, 75 (1992).

27. M. Rauzi, P. Verant, T. Lecuit, P. F. Lenne, Nature and anisotropy of cortical forces orienting Drosophila tissue morphogenesis. Nat Cell Biol 10, 1401– 1410 (2008).

28. N. D. Czerniak, K. Dierkes, A. D’Angelo, J. Colombelli, J. Solon, Patterned Contractile Forces Promote Epidermal Spreading and Regulate Segment Positioning during Drosophila Head Involution. Current Biology 26, 1895–1901 (2016).

29. A. Sumi, P. Hayes, A. D’Angelo, J. Colombelli, G. Salbreux, K. Dierkes, J. Solon, Adherens Junction Length during Tissue Contraction Is Controlled by the Mechanosensitive Activity of Actomyosin and Junctional Recycling. Dev Cell 47, 453–463.e3 (2018).

30. X. Xie, J. A. Fischer, On the roles of the Drosophila KASH domain proteins Msp-300 and Klarsicht. Fly (Austin) 2, 74–81 (2008).

31. G. R. Kidiyoor, Q. Li, G. Bastianello, C. Bruhn, I. Giovannetti, A. Mohamood, G. V. Beznoussenko, A. Mironov, M. Raab, M. Piel, U. Restuccia, V. Matafora, A. Bachi, S. Barozzi, D. Parazzoli, E. Frittoli, A. Palamidessi, T. Panciera, S. Piccolo, G. Scita, P. Maiuri, K. M. Havas, Z. W. Zhou, A. Kumar, J. Bartek, Z. Q. Wang, M. Foiani, ATR is essential for preservation of cell mechanics and nuclear integrity during interstitial migration. Nature Communications 2020 11:1 11, 1–16 (2020).

32. D. Kong, Z. Lv, M. Häring, B. Lin, F. Wolf, J. Großhans, In vivo optochemical control of cell contractility at single-cell resolution. EMBO Rep 20, e47755 (2019).

33. S. Henretta, J. Lammerding, Nuclear envelope proteins, mechanotransduction, and their contribution to breast cancer progression. npj Biological Physics and Mechanics 2025 2:1 2, 1–11 (2025).

34. J. Aureille, V. Buffière-Ribot, B. E. Harvey, C. Boyault, L. Pernet, T. Andersen, G. Bacola, M. Balland, S. Fraboulet, L. Van Landeghem, C. Guilluy, Nuclear envelope deformation controls cell cycle progression in response to mechanical force. EMBO Rep 20 (2019).

35. S. Regot, J. J. Hughey, B. T. Bajar, S. Carrasco, M. W. Covert, High-sensitivity measurements of multiple kinase activities in live single cells. Cell 157, 1724 (2014).

36. A. L. Godeau, M. Marin-Riera, E. Trubuil, S. Rogalla, G. Bengoetxea, L. Backová, T. Pujol, J. Colombelli, J. Sharpe, E. Martin-Blanco, J. Solon, A transient contractile seam promotes epithelial sealing and sequential assembly of body segments. Nature Communications 2025 16:1 16, 1–14 (2025).

37. R. Strick, P. L. Strissel, K. Gavrilov, R. Levi-Setti, Cation-chromatin binding as shown by ion microscopy is essential for the structural integrity of chromosomes. J Cell Biol 155, 899–910 (2001).

38. R. Phengchat, H. Takata, K. Morii, N. Inada, H. Murakoshi, S. Uchiyama, K. Fukui, Calcium ions function as a booster of chromosome condensation. Sci Rep 6, 38281 (2016).

39. H. Cheng, N. Zhang, D. Pati, Cohesin subunit RAD21: From biology to disease. Gene 758, 144966 (2020).

40. Y. Zhu, M. Denholtz, H. Lu, C. Murre, Calcium signaling instructs NIPBL recruitment at active enhancers and promoters via distinct mechanisms to reconstruct genome compartmentalization. Genes Dev 35, 65–81 (2021).

41. D. E. Ingber, N. Wang, D. Stamenović, Tensegrity, cellular biophysics, and the mechanics of living systems. Rep Prog Phys 77, 046603 (2014).

42. D. F. Meaney, B. Morrison, C. D. Bass, The Mechanics of Traumatic Brain Injury: A Review of What We Know and What We Need to Know for Reducing Its Societal Burden. J Biomech Eng 136, 0210081 (2014).

43. B. Rashid, M. Destrade, M. D. Gilchrist, Mechanical characterization of brain tissue in compression at dynamic strain rates. J Mech Behav Biomed Mater 10, 23–38 (2012).

44. K. A. Barbee, Mechanical Cell Injury. Ann N Y Acad Sci 1066, 67–84 (2006).

45. M. Pachitariu, C. Stringer, Cellpose 2.0: how to train your own model. Nature Methods 2022 19:12 19, 1634–1641 (2022).

46. D. Ershov, M. S. Phan, J. W. Pylvänäinen, S. U. Rigaud, L. Le Blanc, A. Charles-Orszag, J. R. W. Conway, R. F. Laine, N. H. Roy, D. Bonazzi, G. Duménil, G. Jacquemet, J. Y. Tinevez, TrackMate 7: integrating state-of-the-art segmentation algorithms into tracking pipelines. Nature Methods 2022 19:7 19, 829–832 (2022).

