## Supplementary figures for "Mechanosensitive and Reversible Chromatin-Lamina Dewetting Triggers Cellular Contraction During Wound Healing"

### Figure S1

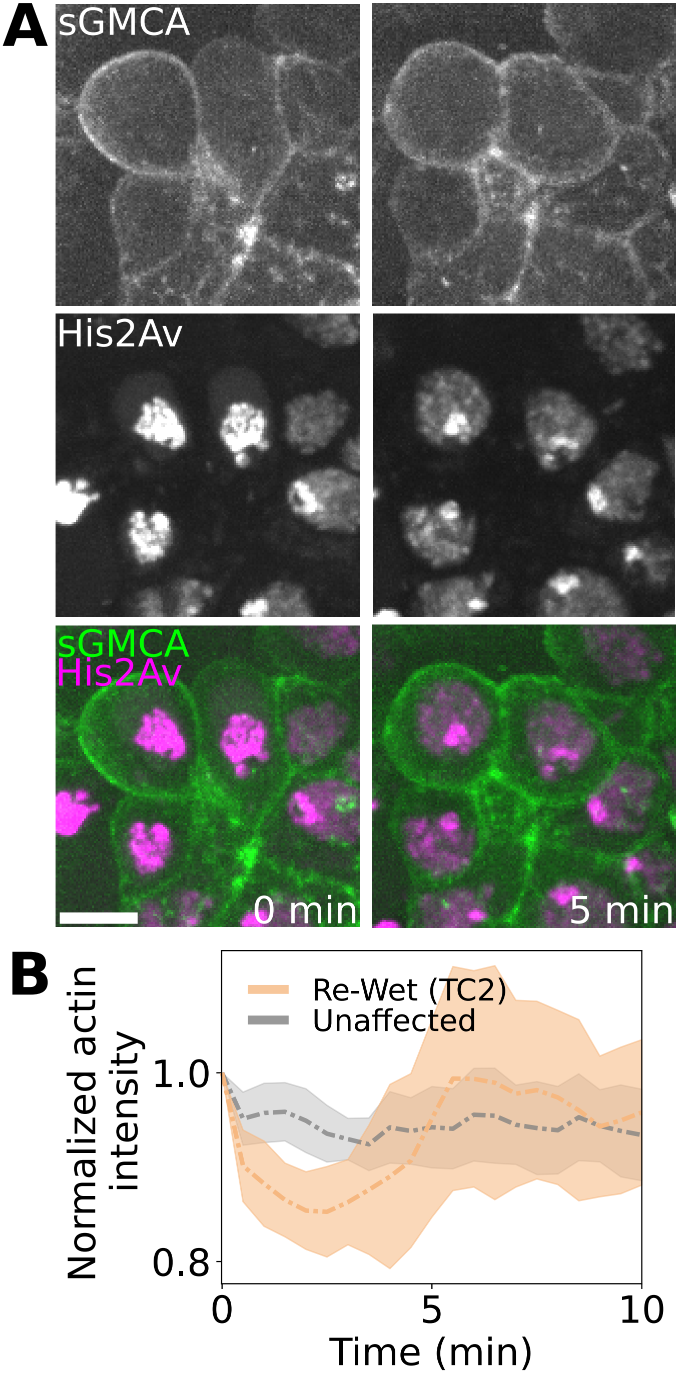
A
sGMCA
His2Av
sGMCA
His2Av
0 min
5 min
B
Re-Wet (TC2)
Unaffected
Normalized actin intensity
1.0
0.8
0
5
10
Time (min)

**Figure S1: Reversible chromatin-lamina dewetting is independent of ultraviolet damage and cortical F-actin behavior in the epidermal cells. (A)** Time-lapse images showing cells undergoing transient chromatin compaction following mechanical wounding (without laser dissection) of the epidermis in embryos expressing sGMCA and His2Av-mRFP. Scale bar 5 μm. **(B)** Graph showing the normalized actin intensity in Re-Wet (TC2) and unaffected cells ($N$ = 5, $n_{TC2}$ = 47, $n_{unaffected}$ = 42, median + mad).

### Figure S2

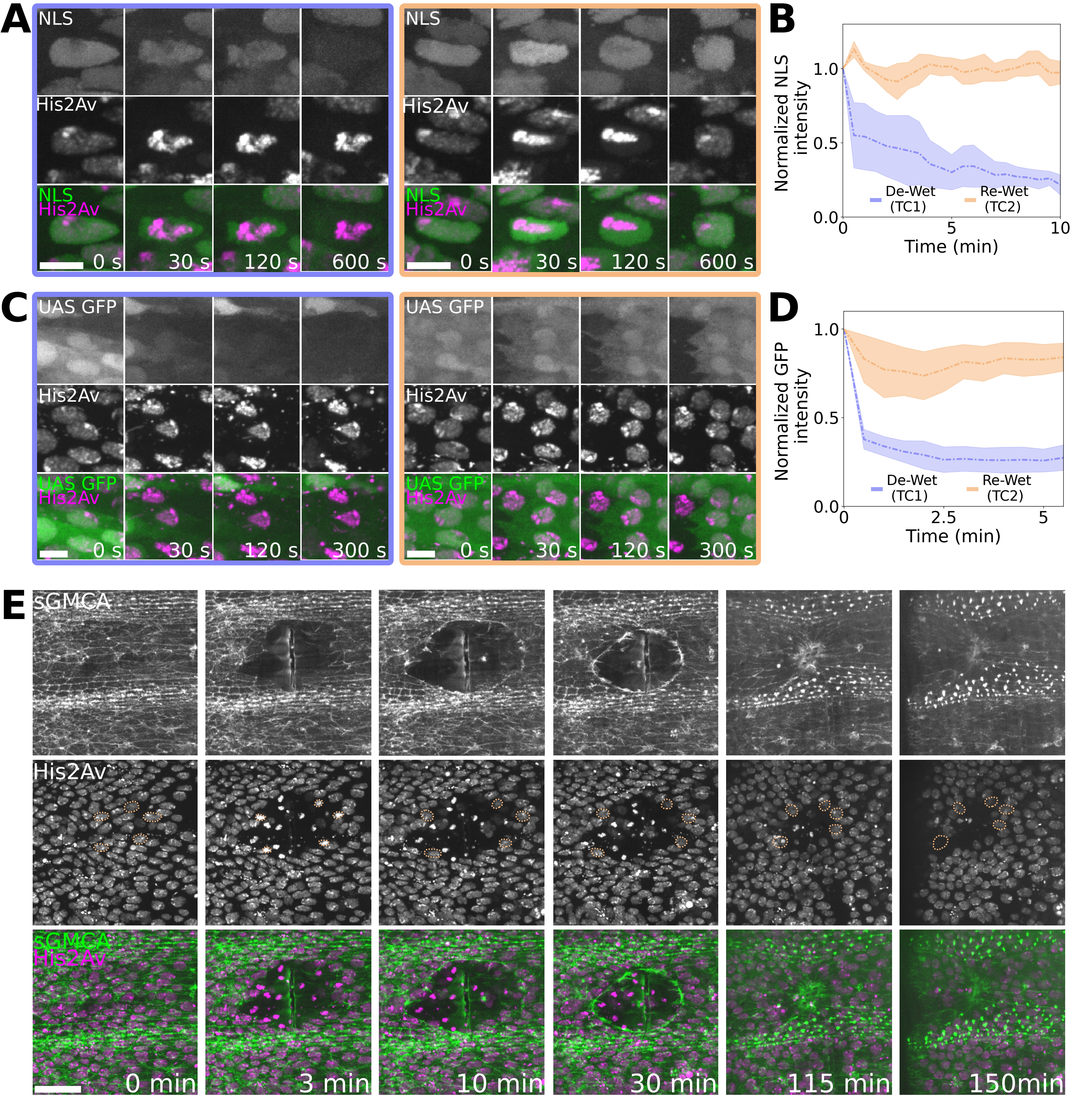
A
NLS
His2Av
NLS
His2Av
0 s
30 s
120 s
600 s
B
Normalized NLS intensity
De-Wet (TC1)
Re-Wet (TC2)
Time (min)
C
UAS GFP
His2Av
UAS GFP
His2Av
0 s
30 s
120 s
300 s
D
Normalized GFP intensity
De-Wet (TC1)
Re-Wet (TC2)
Time (min)
E
sGMCA
His2Av
sGMCA
His2Av
0 min
3 min
10 min
30 min
115 min
150min

**Figure S2: Re-Wet cells maintain compartment integrity during transient chromatin-lamina dewetting. (A)** Time-lapse images showing the decay of NLS signal in the nucleus of a De-Wet (TC1) cell (blue), and a persistence of NLS signal in the nucleus of a Re-Wet (TC2) cell (orange) in embryos expressing NLS GFP and His2Av-mRFP. These data indicate that the De-Wet (TC1) cells undergo nucleoplasmic leakage, whereas Re-Wet (TC2) cells maintain nuclear membrane integrity. Scale bar 5 µm. **(B)** Normalized NLS intensity after wounding in De-Wet (TC1) cells (blue) and Re-Wet (TC2) cells (orange) ($N$ = 6 embryos, $n_{TC1}$ = 17 cells, $n_{TC2}$ = 18 cells, median + mad shown). **(C)** Time-lapse images showing the decay of cytoplasmic GFP signal in a De-Wet (TC1) cell (blue) and persistence in a Re-Wet (TC2) cell (orange). The decay of GFP signal in De-Wet (TC1) cells indicates cytoplasmic leakage consequent of cell damage, while the constant GFP signal in Re-Wet (TC2) cells suggests no noticeable cell damage. Scale bar 5 µm. **(D)** Normalized intensity of GFP over time in De-Wet (TC1) cells (blue) and Re-Wet (TC2) cells (orange). ($N$ = 5, $n_{TC1}$ = 16, $n_{TC2}$ = 16, median + mad shown). **(E)** Time-lapse images showing that the Re-Wet (TC2) cells (orange) remain in the tissue, indistinguishable from other epidermal cells, during healing and post-closure in embryos expressing sGMCA and His2Av-mRFP. Scale bar 20 µm.

### Figure S3

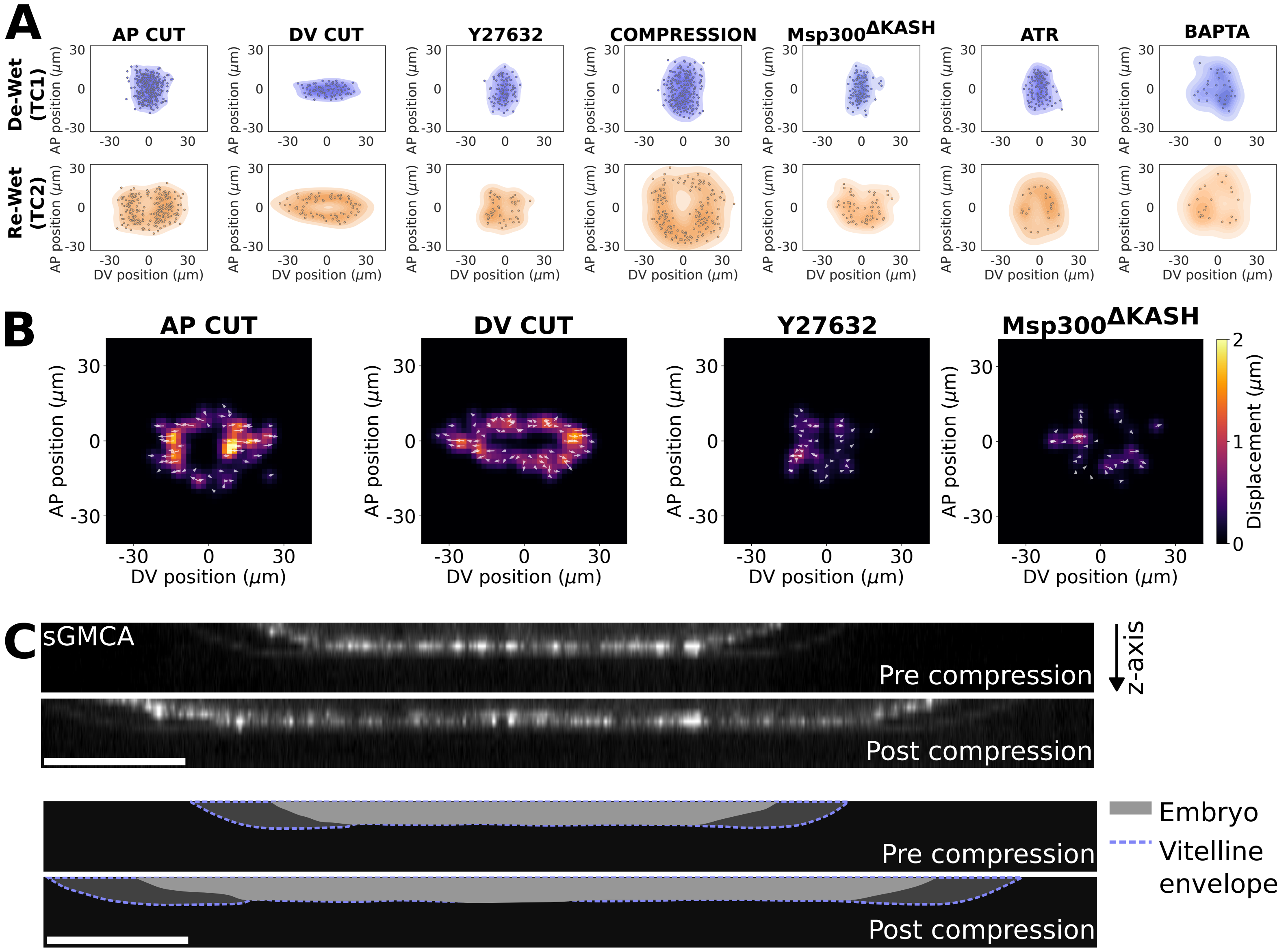
A
AP CUT
DV CUT
Y27632
COMPRESSION
Msp300ΔKASH
ATR
BAPTA
De-Wet (TC1)
Re-Wet (TC2)
AP position (μm)
DV position (μm)
B
AP CUT
DV CUT
Y27632
Msp300ΔKASH
AP position (μm)
DV position (μm)
Displacement (μm)
C
sGMCA
Pre compression
Post compression
z-axis
Embryo
Vitelline envelope
Pre compression
Post compression

**Figure S3: Chromatin re-wetting is mechanosensitive. (A)** Probability distribution maps in the damaged area of De-Wet (TC1) cells and Re-Wet (TC2) cells in the following experimental settings: comparing the wounds induced along the AP axis, along the DV axis, Y27632-injected embryos, compressed embryos, Msp300$^{\Delta KASH}$ mutants, ATR-inhibited embryos, and BAPTA-treated embryos. The graphs relate to Figure 2 B, D, F, Figure 3 A, Figure 4 B, G, and Figure 5 F. **(B)** Deformation field around the wound of Re-Wet (TC2) cells in the following experiments: comparing the wounds induced along the AP axis, along the DV axis, Y27632-injected embryos, and the Msp300$^{\Delta KASH}$ mutants ($N_{AP}$ = 6, $n_{AP}$ = 59, $N_{DV}$ = 76, $n_{DV}$ =6, $N_{Y27}$ = 7, $n_{Y27}$ = 44, $N_{\Delta KASH}$ = 5, $n_{\Delta KASH}$ = 33). **(C)** Images showing the transverse section of an embryo before and after the application of an external force in embryos expressing sGMCA. The surface of the embryo in contact with the coverslip and the amount of epidermis in contact with the vitelline envelope increase during the compression phase of the experiment. The changes are highlighted by a schematic at the bottom. Scale bar 20 µm.

### Figure S4

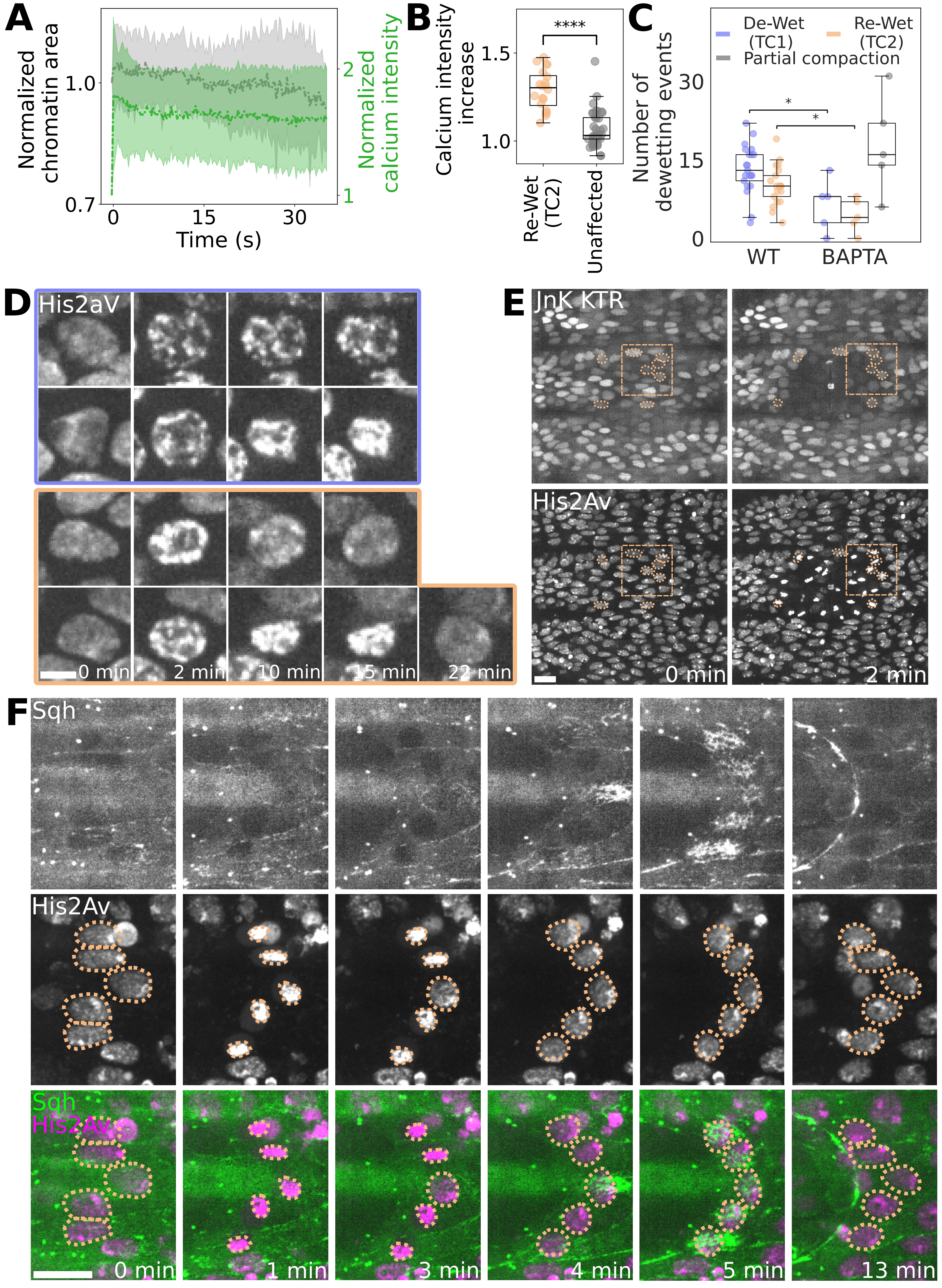
A
Normalized chromatin area
Normalized calcium intensity
1.0
0.7
2
1
0
15
30
Time (s)
B
Calcium intensity increase
1.5
1.0
****
Re-Wet (TC2)
Unaffected
C
De-Wet (TC1)
Re-Wet (TC2)
Partial compaction
Number of dewetting events
30
15
0
*
*
WT
BAPTA
D
His2aV
0 min
2 min
10 min
15 min
22 min
E
JnK KTR
His2Av
0 min
2 min
F
Sqh
His2Av
Sqh
His2Av
0 min
1 min
3 min
4 min
5 min
13 min

**Figure S4: Chromatin compaction is calcium-dependent, and Re-Wetting cells precede wound healing mechanisms. (A)** Graph showing the normalized chromatin area and normalized calcium intensity in unaffected cells close to the wound after the wound was induced ($N$ = 2, $n$ = 10). The following behavior can be observed: rapid increase in calcium intensity during the calcium wave, and a subsequent stabilization of the signal. The onset of time is after the cut was induced, when the fluorescence calcium signal stabilized. **(B)** Graph showing a higher calcium increase in Re-Wet (TC2) cells compared to the unaffected cells in BAPTA-treated embryos. This indicates that cells are able to perform a reversible chromatin-lamina dewetting in these conditions because of remnant calcium ($N$ = 5, $n_{TC2}$ = 20, $n_{unaffected}$ = 38, $p < 0.0001$). **(C)** Time-lapse images showing partial compaction of chromatin in BAPTA-injected embryos expressing His2AV-mRFP in De-Wet (TC1) (blue) and Re-Wet (TC2) cells (orange). In both cases, the chromatin undergoes a partial compaction, with slow kinetics, suggesting that the chromatin does not disassemble from the lamina. In a subpopulation of the nuclei with partially compacted chromatin, the chromatin eventually fully compacts with a strong delay. Scale bar 5 µm. **(D)** Number of dewetting events (TC1 and TC2), and partial compaction events in WT and BAPTA-treated embryos, showing a reduction of TC1 and TC2 cases in BAPTA-injected embryos and the emergence of partial compaction events ($N_{WT}$ = 5, $N_{BAPTA}$ = 5, $p_{TC1}$ = 0.0339, $p_{TC2}$ = 0.0111, two-tailed $t$ test). **(E)** Images of the wound showing that JnK is not activated right after the cut in embryos expressing Jnk KTR and His2AV-mRFP. Re-Wet (TC2) cells are outlined in orange, and the bounding box close-up can be found in the main figure. Scale bar 20 µm. **(F)** Time-lapse images of myosin burst in Re-Wet (TC2) cells after re-wetting occurs and subsequent apical side contraction and recruitment of myosin for actomyosin cable formation in embryos expressing Sqh GFP and His2AV-mRFP. Nuclei are outlined with orange dashed lines. Scale bar 10 µm.

**Description of Additional Supplementary Files**

**Movie S1.**
Tissue retraction after wound generation in an embryo expressing His2Av mRFP. In a subset of the cells, chromatin compacts without re-expansion (TC1); in another subset of cells, chromatin compacts and after a few minutes re-expands. Scale bar 20 μm.

**Movie S2.**
Close-up view of a TC2 cell showing chromatin compaction and re-expansion. Scale bar 5 μm.

**Movie S3.**
Close-up view of a TC2 cell of an embryo expressing Lamin GFP and His2Av mRFP. The chromatin de-wets from the lamina, and consequently, the nucleus becomes spherical. A few minutes later, the chromatin re-wets the lamina, and the nucleus recovers its shape. Scale bar 5 μm.

**Movie S4.**
Wound generation and the affected retraction in an embryo injected with ROCK inhibitor Y27632 and expressing His2Av mRFP. The tissue retraction is smaller than in WT. Scale bar 20 μm.

**Movie S5.**
Compression experiment. An embryo expressing sGMCA and His2Av mRFP is under ectopic pressure during the wound generation. The pressure is released 15 minutes after the wound is generated. The re-wetting in Re-Wet (TC2) cells occurs after pressure is released. Scale bar 20 μm.

**Movie S6.**
Wound generation and the affected retraction in a homozygous $Msp300^{\Delta KASH}$ mutant embryo expressing His2Av mRFP. The tissue retraction is smaller, and we detect fewer cases of re-wetting compared to the Wild Type. Scale bar 20 μm.

**Movie S7.**
Close-up view of a TC2 cell, a TC1 cell, and an unaffected cell in embryos expressing GCaMP8f and His2Av. Depletion of free intracellular calcium occurs in the TC2 cell in the de-wet state, and a burst of calcium concurrently with the re-wetting. Scale bar 5 μm.

**Movie S8.**
Wound generation and affected dynamics in an embryo injected with the permeant calcium chelator BAPTA-AM expressing His2Av mRFP. Chromatin compaction is affected, and we observe a reduction of Re-Wet (TC2) cells. Scale bar 20 μm.

**Movie S9.**
Close-up view of a segment of the wound's leading edge and subsequent start of healing in an embryo expressing sGMCA and His2Av mRFP. Soon after the chromatin-lamina re-wetting, an actomyosin burst occurs in the cell with a following apical surface reduction and onset of actomyosin cable formation. Scale bar 10 μm.
